# AI/ML-Assisted Computational Design and Immunoinformatics Evaluation of a Multi-Epitope Vaccine Targeting Podoplanin in *Glioblastoma Multiforme*

**DOI:** 10.64898/2026.02.18.706629

**Authors:** Gopika Anilkumar, Ranbir Saluja, Aarshit Mittal, Parmi Shah, Shobhan Shah, Prashant S. Kharkar

**Affiliations:** Department of Pharmaceutical Sciences and Technology, Institute of Chemical Technology, Nathalal Parekh Marg, Matunga, Mumbai 400 019. India; Aarth Software Private Limited, Metro Station, 4th Floor, Vasavi MPM Grand, Beside Ameerpet Metro Station, Hyderabad, Telangana, 500073, India; Rasayan Labs Inc., 1340 S De Anza Blvd Ste #208, San Jose, CA 95129, USA

**Keywords:** Glioblastoma multiforme, GBM, Podoplanin, PDPN, Vaccine

## Abstract

Glioblastoma Multiforme (GBM) is one of the most malignant forms of brain tumor in humans, with limited treatment options and poor overall survival rates. In the present study, we employed an in-silico workflow that integrated immunoinformatics and 3D structural modelling tools to design a multi-epitope vaccine against Podoplanin (PDPN), a transmembrane glycoprotein primarily involved in tumor invasion and metastasis. The differential expression of PDPN in tumor versus normal cells was investigated using transcriptomics datasets. Once the overexpression was confirmed, it was designated as a Tumor-Associated Antigen (TAA). B-cell, CTL, and HTL epitopes were predicted and screened for antigenicity, non-allergenicity, and non-toxicity. Selected epitopes were linked with appropriate adjuvant and linker sequences to construct a vaccine candidate. Codon optimization and in silico cloning was conducted to evaluate the construct’s expression in a mammalian expression vector. The 3D structure of the vaccine candidate was modelled, refined, and validated before molecular docking with immune receptors and immune simulation studies. The results indicated that proposed polypeptide, *RasIC*-01v, could be a potential vaccine candidate for highly vigorous and dangerous cancer like GBM. Further experimental and immunological validations would be required to validate the commercial feasibility and development of *RasIC*-01v.

**Graphical Abstract:** 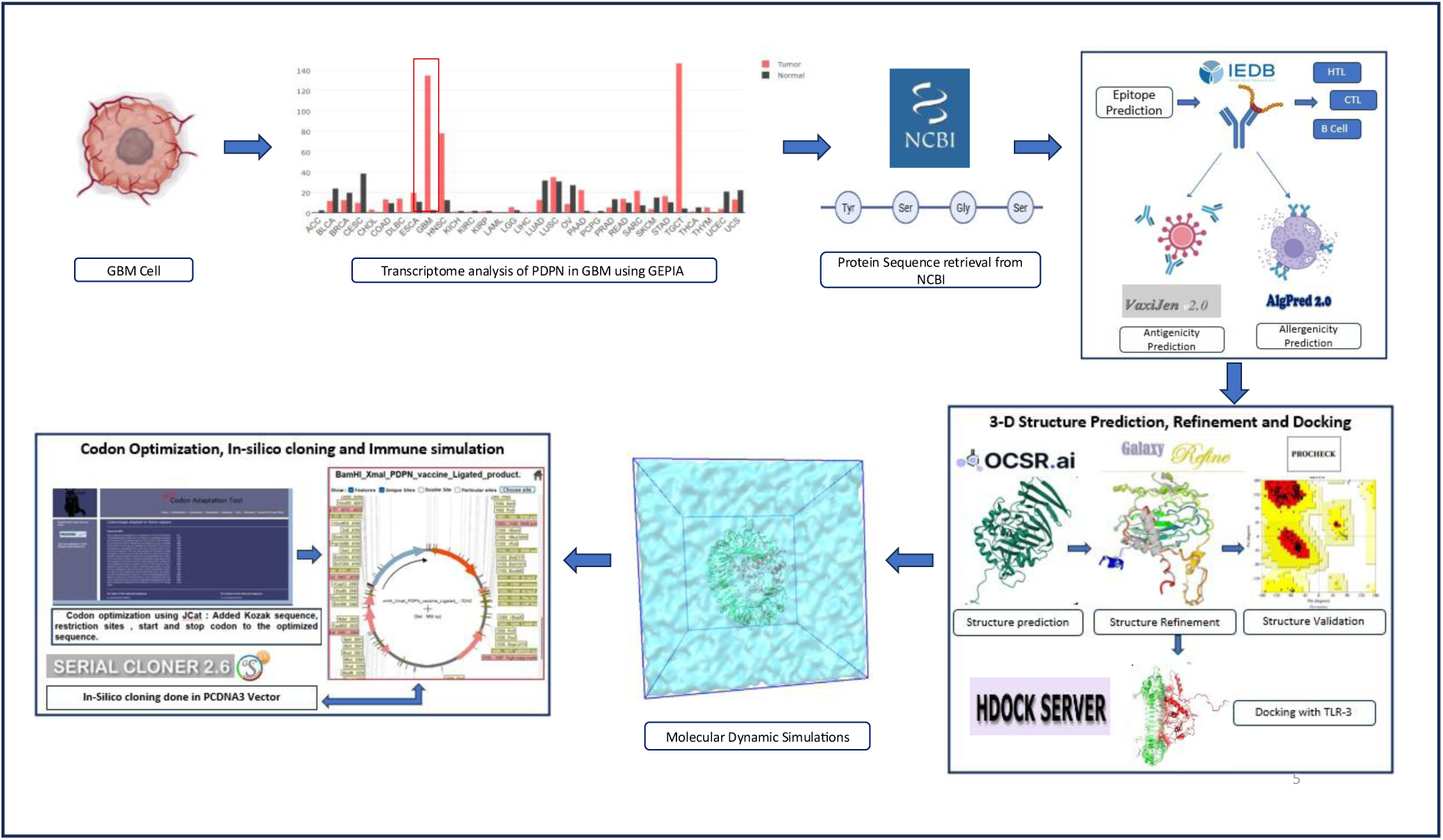

## Introduction

Glioblastoma Multiforme (GBM) is the most malignant forms of astrocytoma, characterised by rapid proliferation, invasiveness and therapeutic resistance. Despite major available treatments like surgical resection, chemotherapy and radiation therapy, the median survival rates are 12-15 months.^1^ The complexity and heterogeneity of GBM and the presence of an immunosuppressive tumour microenvironment limit the effectiveness of conventional therapies. Furthermore, the blood-brain barrier (BBB) restricts the entry of therapeutic molecules and causing insufficient delivery, consequently reducing the overll efficiency of therapeutic agents. One of the methods to bypass this barrier may involve the use of vaccine-induced systemic immunity. It has been proved that in cases of inflammations, increased expression of cell-adhesion molecules like P-selectin along with proinflammatory cytokines and reduced expression of junctional molecules allow increased entry of circulating leukocytes especially T-cells across the BBB.^2^

One of the emerging advances in cancer treatment is immunotherapy, which works by activating the patient’s own immune system to target and destroy the cancer cells. The immune system’s ability to precisely identify and attack these cells helps minimize damage to normal tissues, while also creating a memory of the tumor cells to prevent recurrence or relapse. Checkpoint inhibitors, monoclonal antibodies, CAR-T cell therapy and cancer vaccines are the major modalities of immunotherapy.

Recent advances in computational immunology and AI/ML-based bioinformatics have enabled rapid identification of tumour-associated antigens (TAAs) and epitopes, paving the way for rational vaccine design.^3^ Developing a conventional vaccine takes too much time as it necessitates extensive laboratory testing before it goes into clinical trials. Immunoinformatics driven subunit and multi-epitope vaccine design studies have shown that in silico screening is capable of efficiently identifying conserved, non-toxic, and highly immunogenic epitopes, and simultaneously assessing their stability and receptor-binding affinity using molecular docking and molecular dynamics simulations.^4^ For example, immunoinformatics-driven approaches have enabled the systematic identification of resistance-associated *Mycobacterium tuberculosis* antigens using purely computational pipelines. Whole-genome and proteome data from resistant clinical isolates were screened to identify conserved, surface-exposed virulence proteins, thereby efficiently prioritizing promising vaccine candidates for subsequent experimental validation.^4^ This pre-experimental filtering stage helps to significantly reduce the search space, thus saving time, effort, and resources. Moreover, comprehensive reviews have emphasized that these computational pipelines accelerate vaccine development timelines by providing insights into antigen design, epitope combinations, and immune response prediction across different populations, especially during large-scale vaccine development projects.

Podoplanin (PDPN) also known as AGGRUS, GP36, GP40, Gp38, HT1A-1, OTS8, PA2.26, T1A, T1A-2, T1A2, or TI1A, a mucin-type transmembrane glycoprotein which contains a single transmembrane domain, a heavily glycosylated extracellular domain and a short nine amino acid cytoplasmic tail, is overexpressed in GBM and contributes to tumour cell migration and platelet aggregation.^5^ It is mostly expressed during the neonatal stage and is crucial for the development of multiple organs like lungs, heart and lymphatic system. Given its restricted expression in normal brain tissue, PDPN represents an ideal immunotherapeutic target. Although PDPN is overexpressed in GBM, it is also expressed on endothelial cells and fibroblastic reticular cells in lymphoid organs in adults.^6^ PDPN expression in cancer cells is precarious as it leads to formation of more stem cells. This broader expression limits the use of PDPN in advanced immunotherapy approaches like CAR-T cell therapy, as these methods are highly specific and potent, but they can also lead to the destruction of normal cells alongside tumor cells. This is where subunit vaccines come into play. While they may not trigger as strong an immune response as CAR-T cells, they still can activate immune cells, allowing for tumor destruction with significantly less background cytotoxicity.

Considering the aggressive biology of GBM, its immune-evasive microenvironment, and the limitations imposed by the BBB, there remains a critical unmet need for therapeutic strategies capable of eliciting targeted yet safe antitumour immunity. Podoplanin represents a biologically relevant but therapeutically challenging antigen due to its tumour overexpression coupled with restricted normal tissue distribution. Consequently, a precision-guided immunization strategy that stimulates controlled immune activation without the off-target toxicity associated with highly potent cell-based therapies is particularly attractive. In this context, immunoinformatics-driven vaccine design provides a rational and accelerated framework to identify optimal epitopes and engineer safe antigen constructs prior to experimental validation. Therefore, the present study aims to computationally design and evaluate a multi-epitope subunit vaccine targeting PDPN for GBM, with the objective of generating a selective, immunogenic, and translationally viable therapeutic candidate for this otherwise fatal malignancy.

## Materials and Methods

### Computational Resources and Analysis Methods

Tsarget identification, epitope selection, multi-epitope vaccine construction, structure analysis, refinement, validation and molecular docking were performed using open-source web based tools. 3D Structure prediction was performed using the *Structure Prediction* module (ProteinLab.ai) available in OCSR^TM^.ai, as implemented in Rasayan Lab Inc.’s integrated Drug Discovery and Biologics Suite (www.rasayan.ai), a proprietary AI-based software. Molecular Dynamics (MD) simulations of the predicted vaccine candidate (*Ras*IC-01v) structure were performed using a high-performance computing (HPC) environment enabled with GPU support powered by OCSR^TM^.ai. On similar lines, MD simulations for the docked TLR3-*Ras*IC-01v complex were performed on an in-house Linux HPC server enabled with a 36-core machine with 128 GB CPU and 24 GB A5000 GPU, running CentOS 7 operating system. Table 1 lists the software and tools used for the complete computational workflow for the development of *Ras*IC-01v.

**Table 1.** List of software and tools used for the present work.

| Category | Software and Tools |
| --- | --- |
| Target Identification | GEPIA (Gene Expression Profiling Interactive Analysis) (http://gepia.cancer-pku.cn/) |
| Epitope prediction | NetMHCpan EL 4.1 web server (IEDB)<br>MHC II prediction server (version 2.22) (IEDB)<br>CLBtope (https://webs.iitd.edu.in/raghava/clbtope/index.php) server |
| Antigenicity prediction | VaxiJen v2.0 (http://www.ddg-pharmfac.net/vaxijen/VaxiJen/VaxiJen.html) |
| Toxicity and allergenicity prediction | ToxinPred (https://webs.iitd.edu.in/raghava/toxin-pred/multi_submit.php) and AlgPred 2.0 (https://webs.iitd.edu.in/raghava/algpred2/) |
| Secondary structure prediction | Chau and Fassman Secondary Structure Prediction Server(https://www.biogem.org/tool/chou-fasman/index.php) |
| 3D structure prediction | Structure prediction module (ProteinLab) in OCSR<sup>TM</sup>.ai (https://ocsr.ai/) |
| 3D structure refinement | GalaxyRefine server (http://galaxy.seoklab.org/cgi-bin/submit.cgi?type=REFINE) |
| 3D structure validation | PROCHECK(https://saves.mbi.ucla.edu/) and Prosa-Web(https://prosa.services.came.sbg.ac.at/prosa.php) |
| Molecular docking | HDOCK webserver (http://hdock.phys.hust.edu.cn/) |
| Molecular dynamics simulations | GROMACS 2024.3 and GROMACS 2025.3 |
| Generation of simulation input files | CHARMM GUI server (https://www.charmm-gui.org/) |
| Binding free energy | gmx_MMPBSA v1.6.4 by Valdes tresanco (https://valdes-tresanco-ms.github.io/gmx_MMPBSA/) |
| Structure visualisation and simulation movie generation | PyMOL<sup>TM</sup> v3.1.6.1 (educational version)<br>VMD v1.9.3 |
| Plotting graphs | QtGrace v0.2.6 (https://sourceforge.net/projects/qtgrace/) |
| Codon optimization and in silico cloning | VectorBuilder (https://en.vectorbuilder.com/) server. SerialCloner (http://serialbasics.free.fr/Serial_Cloner.html) |
| Immune simulation | C-ImmSim (https://kraken.iac.rm.cnr.it/C-IMMSIM/index.php) |

### Target Identification

In the present study, GBM was chosen as a disease model and different Tumor Associated Antigens’ (TAAs) comparative expression was analysed using GEPIA tool (http://gepia.cancer-pku.cn/). GEPIA (Gene Expression Profiling Interactive Analysis) is a web-based platform designed for large-scale transcriptomic analysis by integrating RNA-sequencing data from TCGA and GTEx projects. The processed expression data are stored in a MySQL relational database, enabling efficient querying and real-time analysis.^7^ The targets whose expression is very minute in healthy tissues compared to tumor are selected so as to reduce off-tumor toxicity. The expression pattern was double confirmed using literature survey. Shortlisted target candidates were SOX11, SOX 2, Survivin/BIRC5 and PDPN. Of these, SOX 11 and SOX 2 are both transcription factors that belong to the SOX family which binds to the consensus sequence 5′-(A/T)(A/T)CAA(A/T)G-3′ in the minor groove of DNA and helps in adult neurogenesis.^8^ Survivin/BIRC5 is highly expressed in normal blood cells. Hence, these potential targets were ruled out. In case of PDPN, it is highly expressed in neonatal cells only and expression is restricted in adult brain cells while overexpressed in GBM cells, which made it an ideal immunotherapy vaccine candidate.

### Epitope Prediction

#### Cytotoxic T Lymphocyte (CTL) Epitope

To identify potential CTL epitopes within the selected PDPN antigen sequence, predictions were carried out using the NetMHCpan EL 4.1 web server (available at <u>IEDB</u>).^9^ This platform employs artificial neural network (ANN) models combined with position-specific scoring matrices to estimate peptide–MHC class I binding affinity. To include 97% population coverage, 27 reference HLA alleles suggested by IEDB were used. The CTL epitopes with IC_50_ values <100 were considered and their antigenicity as well as allergenicity were checked. CTL epitopes help in binding to MHC I and induce cellular cytotoxicity to the antigen.

#### Helper T Lymphocyte (HTL) Epitope

Antigenic epitopes are presented by Antigen Presenting Cells (APCs) as epitope-MHC II complex to helper T-cells. MHC class II-restricted epitopes were predicted using the IEDB MHC II prediction server (version 2.22) (<u>IEDB</u>). The analysis was performed with parameters set for *Homo sapiens*, HLA-DR alleles, and a peptide length of 15 amino acids. A reference panel of seven representative HLA-DR alleles was utilized to ensure broad population coverage. Epitopes showing minimum percentile, IC_50_ and adjusted ranks were considered to have high binding affinity and were selected for further screening.

#### Linear B-cell Lymphocyte Epitope

B-Cell epitopes are inevitable for humoral immune response which in turn induces antibody production while the epitope of antigen meets with a B-cell receptor. CLBtope (https://webs.iiitd.edu.in/raghava/clbtope/index.php)^10^ server was used to predict B cell epitopes. It employs a hybrid framework that integrates a Random Forest–based machine learning classifier for discriminating B-cell epitopes from non-epitopic peptides, together with a BLAST-based sequence similarity approach that identifies peptides homologous to experimentally validated B-cell epitopes. The server performs three consequent jobs before giving out results; Predict, Scan and Design. In Predict, the software allows us to input possible candidates and it returns the possibility of them being B-cell epitope as a result and then the design part generates mutants from the peptides input and then searches for the probability of the mutant being a B-cell epitope. The final part scan can be mostly conducted by Motif Scan (MERCI) and BLAST scan. Variable epitope length function was chosen for the prediction instead of fixed length (20-mer).

### Antigenicity Prediction

To evaluate the immune-stimulating potential of the obtained epitopes, VaxiJen v2.0 (http://www.ddg-pharmfac.net/vaxijen/VaxiJen/VaxiJen.html)^11^ was employed for the antigenicity assessment. Unlike traditional alignment-based tools that depend on sequence similarity, VaxiJen applies an auto- and cross-covariance transformation method to analyse the physicochemical properties of the amino acids. In this approach, each amino acid in the sequence is represented by numerical values corresponding to properties such as hydrophobicity, polarity, and molecular size. These values are then mathematically transformed into a uniform vector that captures overall patterns and correlations within the sequence. The resulting vector is processed through a discriminant function trained on known datasets of antigens and non-antigens, allowing the tool to classify new sequences according to their likelihood of being antigenic. This makes VaxiJen particularly useful for identifying potential vaccine candidates even when no close homologs exist in the database. In the present study, peptides or proteins showing a prediction score of 0.5 or higher were classified as probable antigens and were shortlisted for further development of the multi-epitope vaccine construct.

### Allergenicity Prediction

To ensure safety and biocompatibility, the selected antigenic peptides were analyzed for toxicity and allergenicity using ToxinPred (https://webs.iiitd.edu.in/-raghava/toxinpred/multi_submit.php)^12^ and AlgPred 2.0 (https://webs.iiitd.edu.in/raghava/-al-gpred2/).^13^ ToxinPred classifies peptides as toxic or non-toxic based on sequence-derived features integrated with ML models, enabling identification of safe epitope candidates. AlgPred 2.0 combines support vector machine algorithms, BLAST-based similarity search, motif analysis (MEME/MAST), and IgE epitope mapping to screen for allergenic determinants. Peptides predicted to be non-toxic and non-allergenic were retained for final vaccine assembly.

### Construction of Multi-epitope Vaccine Candidate Sequence

Epitopes demonstrating strong antigenicity, immunogenicity, and confirmed non-toxic and non-allergenic profiles were selected for inclusion in the final multi-epitope vaccine construct. The shortlisted CTL, HTL, and B-cell epitopes were derived from predictions using IEDB and CLBTope servers, respectively. To maintain proper spacing and enhance independent epitope presentation, AAY linkers were used to connect CTL epitopes for proper proteasomal cleavage, while GPGPG linkers for flexible separation were inserted between HTL and B-cell epitopes. To augment the overall immune stimulation, the Hp91 peptide adjuvant (sequence: DPNAPKRPPSAFFLFCSE), a fragment derived from the HMGB1 B-box domain, was integrated at the N-terminus of the construct using a rigid EAAAK linker. Hp91 has been reported to activate dendritic cells via TLR-dependent signalling, inducing secretion of cytokines such as IL-12 and type I interferons, thereby promoting both cellular and humoral immune responses.^14^ This combination of adjuvant and epitope assembly was designed to maximize the vaccine’s immunogenic potency against GBM.

### Secondary and Tertiary Structure Prediction, Refinement and Validation

Secondary structure prediction was done using Chau and Fassman Secondary Structure Prediction Server (https://www.biogem.org/tool/chou-fasman/index.php), which predicts helix, sheet and turn-forming regions based on empirical residue propensities.^15^ The full amino acid sequence of the vaccine construct, *Ras*IC-01v, was submitted to the server, and the distribution of helices, β-sheets, and random coils was recorded. The 3D structure of *Ras*IC-01v was predicted using the ProteinLab.ai module of OCSR^TM^.ai. Prediction involves steps like deriving atomic coordinates from the primary amino acid sequence, optimising structural geometry and generating confidence metrics. The resulting model was then subjected to stereochemical validation (Ramachandran plot, bond-angle checks) prior to downstream docking. The preliminary 3D model of *Ras*IC-01v was refined using the GalaxyRefine server (http://galaxy.seoklab.org/cgi-bin/submit.cgi?type=REFINE).^16^ This tool employs a multi-step refinement protocol, validated through CASP10 experiments, which integrates side-chain repacking and molecular dynamics–based relaxation to correct local geometry and enhance overall structural quality. The procedure improves both global backbone accuracy and local stereochemical parameters, producing a more reliable structural model for downstream analysis. The predicted 3D structure, when put in GalaxyRefine, gave five models, of which the ideal one was selected mainly on the basis of RMSD and Rama favour score.

Validation of the tertiary structure was then carried out using PROCHECK (https://saves.mbi.ucla.edu/)^17^ and Prosa-Web (https://prosa.services.came.sbg.ac.at/-prosa.php)^18^ to ensure model reliability and detect potential structural inconsistencies. The ProSA-web validation yielded a Z-score of –3.67 for the 293 amino acid vaccine construct, placing it within the range of experimentally resolved protein structures of comparable length. This is consistent with the predominantly disordered nature of multi-epitope vaccines.

### Molecular Docking and Dynamics Simulations. GBM vaccine candidate, RasIC-01v

The predicted 3D structure of *Ras*IC-01v was used as an input for MD simulation using GROMACS 2025.3.^19^ Total seven cycles of NVT, NPT equilibration has been incorporated. NVT (100 ps) and NPT (step 1.1) equilibration were carried out with position restraints of 1000 KJ/mol on the backbone atoms. The restraints were gradually reduced from 1000 to 0 KJ/mol (6 NPT cycles) to ensure efficient relaxation and removal of structural clashes for a period of 60 ns wherein each cycle lasted for 10 ns using different pressure coupling barostats. The first NPT cycle was conducted using Berendsen barostat and further steps were performed using C-rescale barostat. Thereafter, a 100 ns simulation was run and the last frame was extracted for further analysis.

### Docking of RasIC-01v with TLR3 Receptor

The structural coordinates extracted from the last frame of the solo simulation of *Ras*IC-01v was used for docking analysis with toll-like receptor 3 (TLR3). The structure of TLR3 (PDB ID: 1ZIW) is an ectodomain structure and was fed into HDOCK webserver^20^ along with *Ras*IC-01v (last frame from 100 ns simulation as described previously). Template-free docking mode was selected and the DNA-binding region of TLR3 was specified for docking. HDOCK webserver was specifically chosen as to preserve the Glycans bonded to TLR3 structure. His30, His60, His108, His539, Asn541 were chosen as the binding site residues. The best ranked docked structure with highest HDOCK score was selected for molecular dynamics simulation.

### MD Simulations of RasIC-01v-TLR3 Complex

The docked complex was submitted to Glycan reader module in CHARMM GUI server, to make input files for simulation. The Glycan reader module was specifically chosen to handle the glycan modifications on TLR3 which require careful assignment of dihedral parameters. Molecular dynamics simulation was performed using GROMACS 2024.3. After the initial energy minimization using steepest descent protocol, seven cycles of NVT, NPT equilibration were performed. Glycan reader module specifically assigns dihedral constraints for the glycan molecules, which are gradually reduced to accurately sample their ring structure. The same equilibration protocol (*Ras*IC-01v solo simulation) was used where NVT (100 ps) and NPT (step 1.1) equilibration were carried out with position restraints of 1000 kJ/mol on the backbone atoms. The restraints were gradually reduced from 1000 to 0 kJ/mol (six NPT cycles) for a period of 60 ns wherein each cycle lasted for 10 ns using different pressure barostats. The first NPT cycle was conducted using Berendsen barostat and further steps were performed using C-rescale barostat. Thereafter, a 100 ns simulation was run. MMGBSA (Binding free energy using gmx_MMPBSA v1.6.4 by Valdex tresanco)^21^, RMSD, RMSF, SASA, radius of gyration were used as parameters for post-simulation analysis.

### Visualisation Analysis

PyMOL ™ v3.1.6.1^22^ (educational version) (The PyMOL Molecular Graphics System, Version 3.1.6.1 Schrödinger, LLC.), VMD v1.9.3^23^ was utilised to view all the structures extracted post-simulation. Secondary structure prediction was done using Chau and Fassman Secondary Structure Prediction Server (https://www.biogem.org/tool/chou-fas-man/index.php) for both the simulations. QtGrace v0.2.6 (https://sourceforge.net/projects/qtgrace/) was utilised for plotting of all graphs. Script for interface residue analysis was acquired from PyMOL Wiki(https://www.pymolwiki.org/index.php/Main_Page).

### Codon Optimization and In Silico Cloning

Given the codon degeneracy found in various organisms resulting from their specific codon usage bias, it was crucial to perform codon optimization prior to in-silico cloning. For recombinant expression in mammalian cells, the vaccine coding sequence was optimized according to the *Homo sapiens* codon usage using the VectorBuilder (https://en.vectorbuilder.com/) server.^24^ The optimized codon was used for in-silico cloning using SerialCloner (http://serialbasics.free.fr/Serial_Cloner.html) software. Before going for in-silico cloning, primers were designed to amplify the specific vaccine construct. Primer designing included addition of Recognition sequences for HindIII Kozak sequence and start codon in forward primer and EcoRI recognition sequence along with stop codon in reverse primer. The amplified construct was cloned into pcDNA3 mammalian expression vector usually used in CHO cells.^25^

### Immune Simulation Analysis

The immune profile of the designed construct was evaluated using the C-ImmSim^26^ simulation platform. The protein sequence was provided as input, and the simulation was executed under standard configuration without parameter manual adjustment. The server integrates a position-specific scoring method with an agent-based framework to emulate mammalian immune behaviour, including antigen uptake, T-cell priming and B-cell maturation. The model reports antibody titres, T-cell subset dynamics, cytokine release patterns and the establishment of long-term immune memory across repeated antigen exposures.

## Results and Discussion

### Target Identification and Sequence Retrieval

The comparative expression analysis of PDPN target was done using GEPIA. A remarkable tumor-specific upregulation pattern was discovered through comparative transcriptomic profiling of PDPN expression across 33 TCGA cancer types. Normal tissues from corresponding GTEx datasets showed consistently low baseline PDPN expression (Transcripts per Million, TPM <20 across nearly all tissues). On the other hand, PDPN levels were noticeably elevated in a number of tumor cohorts, with tumor samples clearly clustering above normal. The GBM, head and neck squamous cell carcinoma (HNSC), lung squamous cell carcinoma (LUSC), cervical squamous cell carcinoma (CESC), esophageal carcinoma (ESCA), pancreatic adenocarcinoma (PAAD), thyroid carcinoma (THCA), testicular germ cell tumors (TGCT), and skin cutaneous melanoma (SKCM) all showed marked overexpression. Tumor TPM values for these cancers often exceeded 200–600, with some outliers surpassing 1000 TPM, indicating highly-active PDPN transcription in individual tumor samples (Figure 1). Importantly, the robust expression in GBM, highlights PDPN as a viable immunogenic candidate for vaccine-based therapeutic strategies. Elevated PDPN expression in tumor tissue compared with minimal expression in normal brain provides a favourable tumor–normal differential, reducing potential off-target effects.

**Figure 1.**
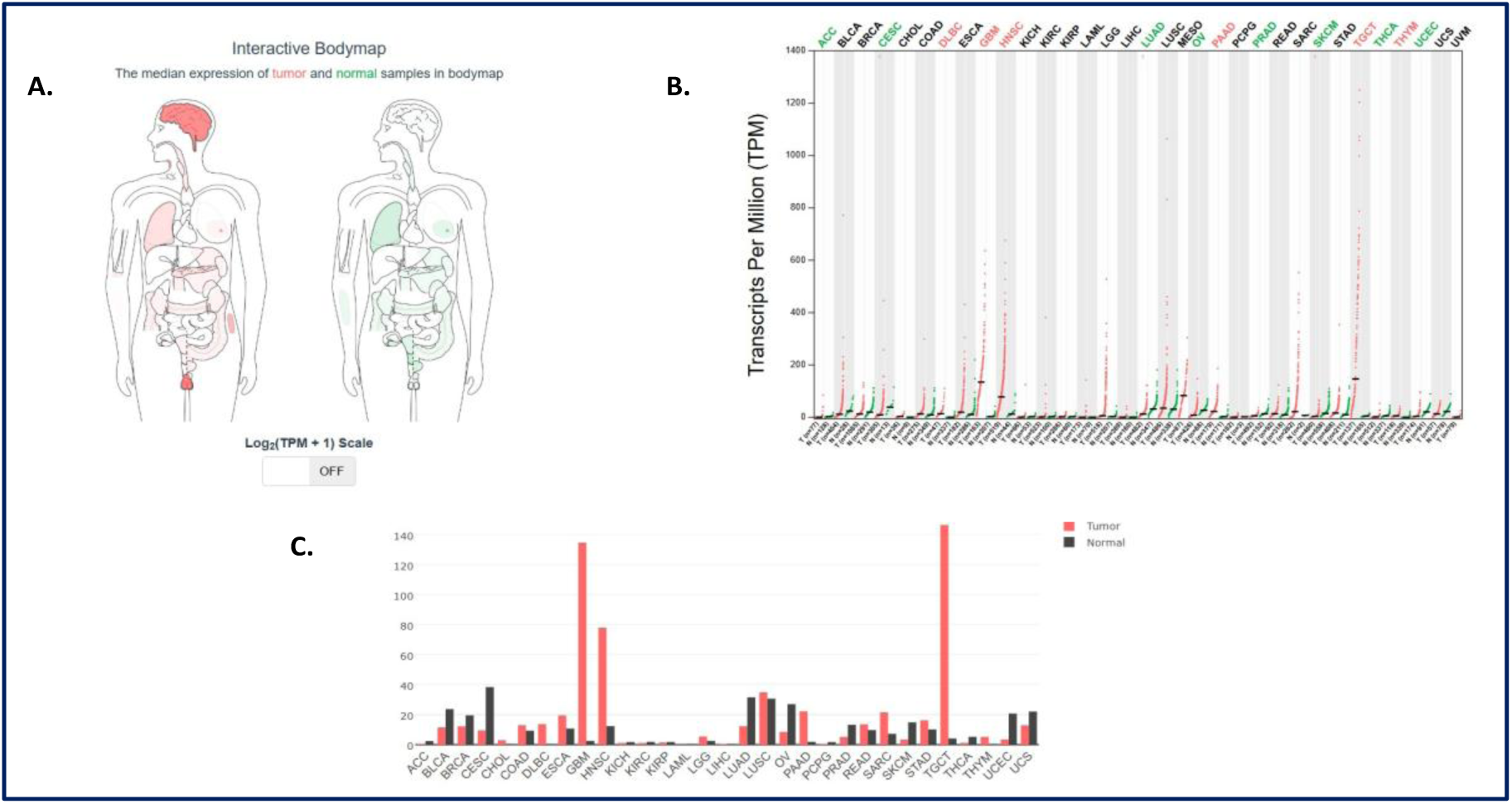
Pan-cancer expression profile of PDPN. A. Interactive bodymap showing median expression of tumor and normal samples of PDPN, B. Scatter plot of TPM values across 33 TCGA cancer types demonstrates markedly higher PDPN expression in several tumors, including GBM, HNSC, LUSC, ESCA and CESC, C. Comparative bar graph showing tumour versus normal tissue expression across cancer types, confirming consistently higher PDPN expression in several tumour cohorts, particularly GBM, supporting its relevance as a potential tumour-associated antigen and therapeutic target.

### HTL, CTL and B-cell Epitope Prediction

A total of 91 HTL (Table 2) and 106 CTL (Table 2) epitopes were selected through IEDB server mainly based on their IC_50_ values being set below 100 and their low percentile score. A total of 75 B-cell epitopes (Table 4) were predicted by using CLBTope server whose threshold hybrid score was above 0.55, from which a total of 32 were selected for further screening based on their higher hybrid score. All these epitopes then went through antigenicity as well as allergenicity screening before being selected for the vaccine construct making.

**Table 2.** HTL Epitopes.

| Allele | Seq. No. | Start | End | Length | Peptide | IC<sub>50</sub> | Percentile Rank |
| --- | --- | --- | --- | --- | --- | --- | --- |
| HLA-DQA1*05:01 | 1 | 15 | 23 | 37 | ARA-PSLSGAPAPTPP | 20.7 | 0.54 |
| HLA-DPA1*01:03 | 1 | 15 | 49 | 63 | LWPRCFLIFPQL-RIL | 24.4 | 0.53 |
| HLA-DQA1*05:01 | 1 | 15 | 22 | 36 | EARA-PSLSGAPAPTP | 25.3 | 0.86 |
| HLA-DQA1*05:01 | 1 | 15 | 110 | 124 | TTGLEGGVAMP-GAED | 37.2 | 1.2 |
| HLA-DPA1*01:03 | 1 | 15 | 46 | 60 | RLGLWPRCFLIFPQL | 40.9 | 1.4 |

**Table 3.** CTL Epitopes.

| Allele | Seq. No. | Start | End | Length | Peptide | IC50 | Percentile Rank |
| --- | --- | --- | --- | --- | --- | --- | --- |
| HLA-A*68:02 | 1 | 109 | 117 | 9 | ETTGLEGGV | 6.67 | 0.07 |
| HLA-A*31:01 | 1 | 44 | 52 | 9 | SSRLGLWPR | 10.56 | 0.09 |
| HLA-A*31:01 | 1 | 43 | 52 | 10 | RSSRLGLWPR | 11.12 | 0.1 |
| HLA-A*02:03 | 1 | 196 | 205 | 10 | TVEKDGLSTV | 27.39 | 0.45 |
| HLA-A*02:03 | 1 | 75 | 83 | 9 | GTMWKVSAL | 29.34 | 0.48 |
| HLA-B*07:02 | 1 | 24 | 33 | 10 | RAPSLSGAPA | 31.07 | 0.12 |
| HLA-A*68:02 | 1 | 108 | 117 | 10 | TETTGLEGGV | 57.23 | 0.41 |
| HLA-A*68:02 | 1 | 181 | 189 | 9 | ATSHSTEKV | 93.59 | 0.57 |

**Table 4.**
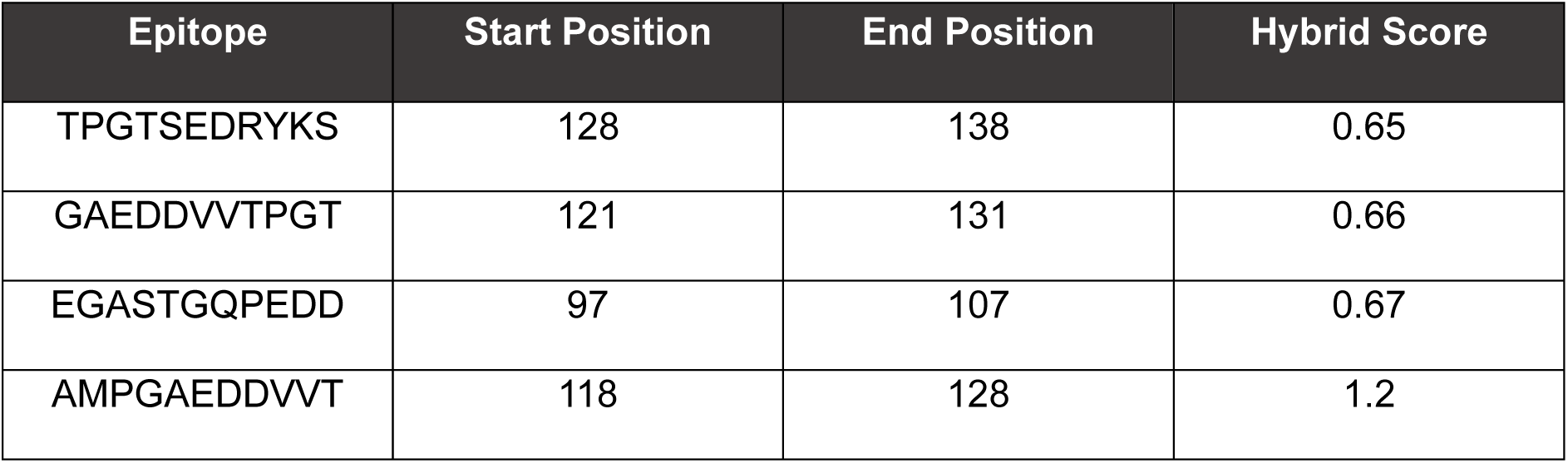
B-cell Epitopes.

**Table 5.**
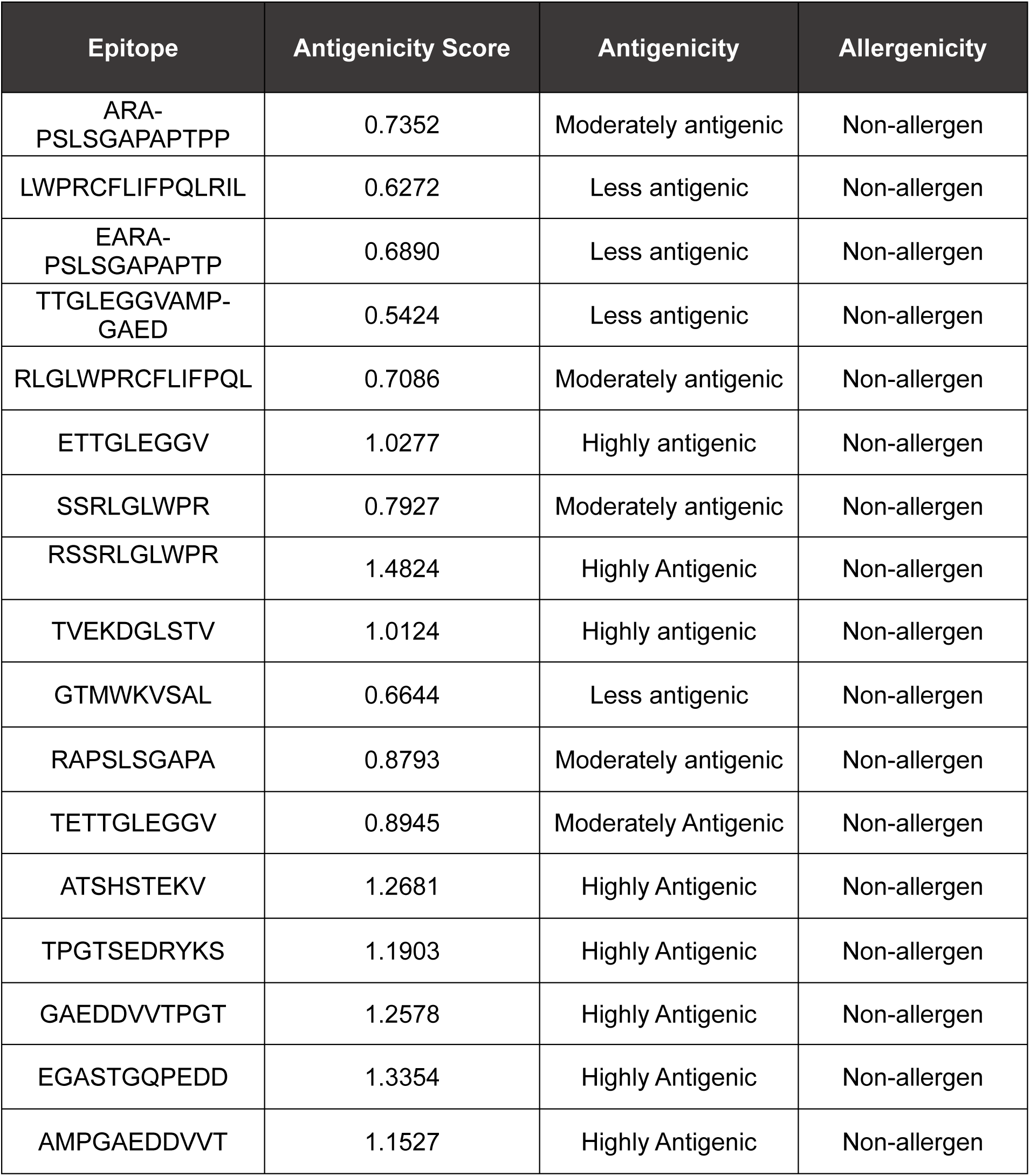
Antigenicity and allergenicity characteristics of the selected epitopes.

### Antigenicity and Allergenicity Prediction

Antigenicity and allergenicity of the selected epitopes of HTL, CTL and B-cell epitopes were predicted (Table 4). Five HTL, eight CTL and Four B-cell epitopes were shortlisted for the construction of vaccine sequence.

### Construction of Multi-epitope Vaccine Sequence, *Ras*IC-01v

Five HTL, eight CTL and four B-cell epitopes were nominated to form a multi-epitope vaccine construct, *Ras*IC-01v, along with Hp-91 peptide as an adjuvant. The schematic representation of the vaccine construct is given below (Figure 2).

**Figure 2.**
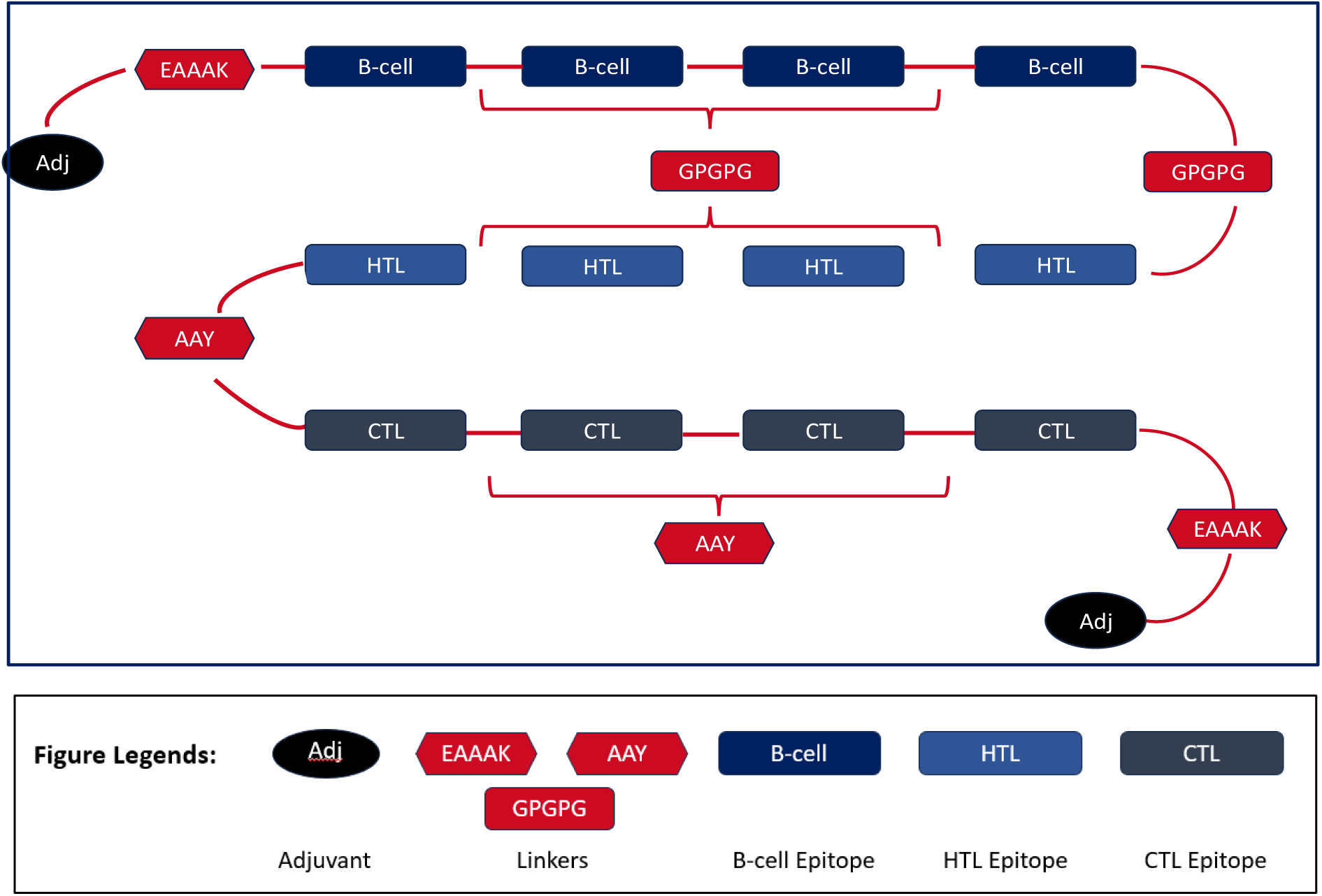
Schematic representation of the final multi-epitope vaccine candidate, *Ras*IC-01v

The 293 amino acid long multi-epitope peptide sequence containing adjuvant at both N and C termini was linked using EEEAK linker. B-cell epitopes and HTL epitopes were linked using GPGPG linkers while CTL epitopes are linked using AAY linkers.

### Secondary Structure Prediction of *Ras*IC-01v

The Chou-Fasman prediction indicated a secondary structure composition of 44% α-helical content, 27% β-sheet architecture and 16% turns, with the remainder forming coils (Figure 3). Such distribution is consistent with a semi-ordered protein fold in which flexible regions may enhance the accessibility of antigenic sur-faces.

**Figure 3.**
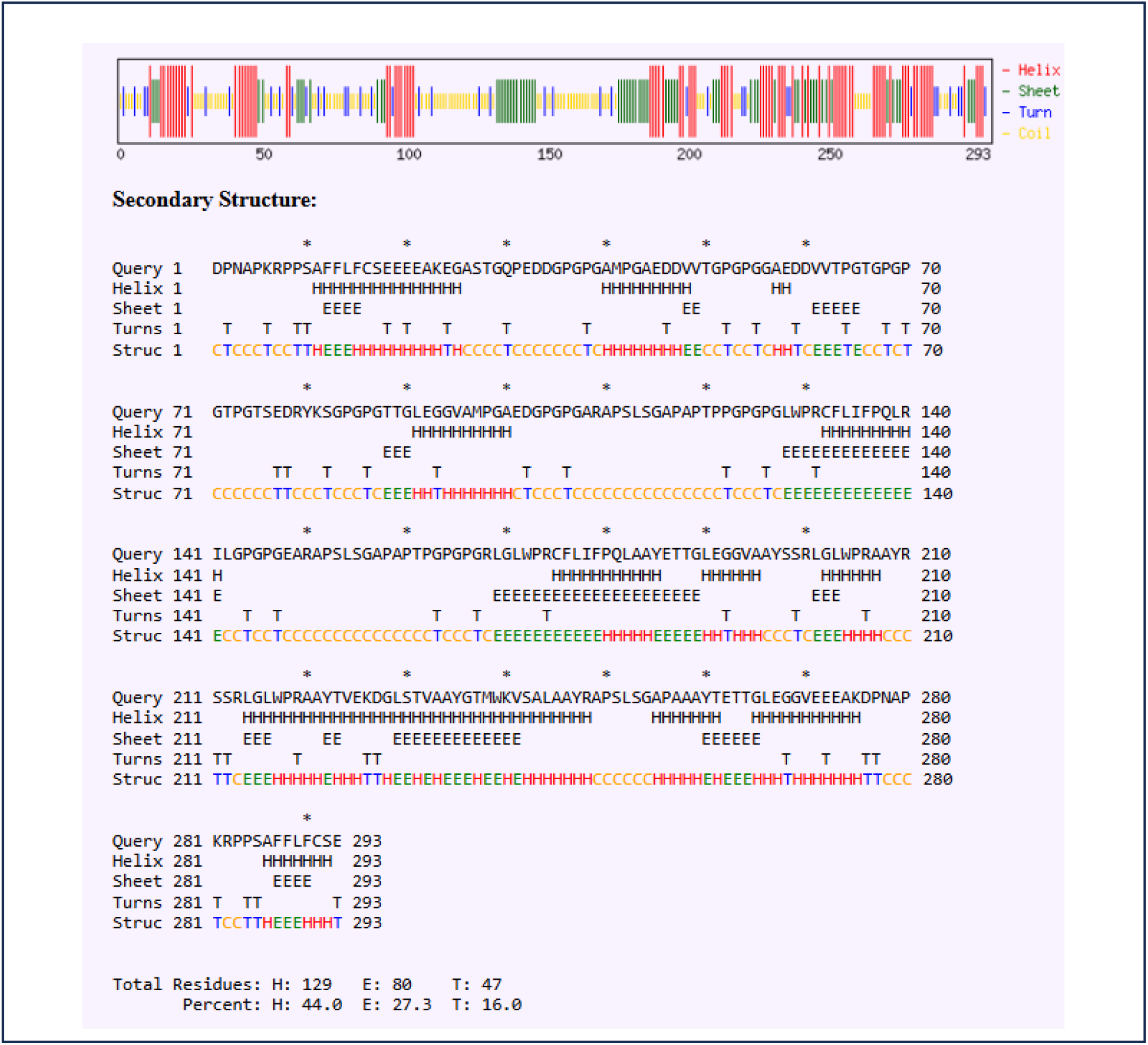
Secondary structure of *Ras*IC-01v predicted using Chau and Fassman server showing the distribution of α-helices, β-strands, turns and coils.

### Tertiary Structure Prediction and Refinement

The 3D structure of the vaccine construct was predicted using Boltz-2 structure prediction framework in OCSR™.ai platform, which gave a confidence score of 26.5% and pLDDT score of 0.303 (Figure 4). The construct was found to be predominantly flexible, a characteristic feature commonly observed in multi-epitope vaccine designs. The predicted 3D structure had 137.6 clash score which was high and hence the structure needed to be refined. Refining of the 3D structure was done using GalaxyRefine (https://galaxy.seoklab.org/cgi-bin/submit.cgi?type=REFINE) and it gave five structural outputs, of which one was selected for further steps like docking and simulation. The selected structure had a reduced clash score of 31.6 with only one poor rotamer and a 98.6% rama favour score (Figure 5).

**Figure 4.**
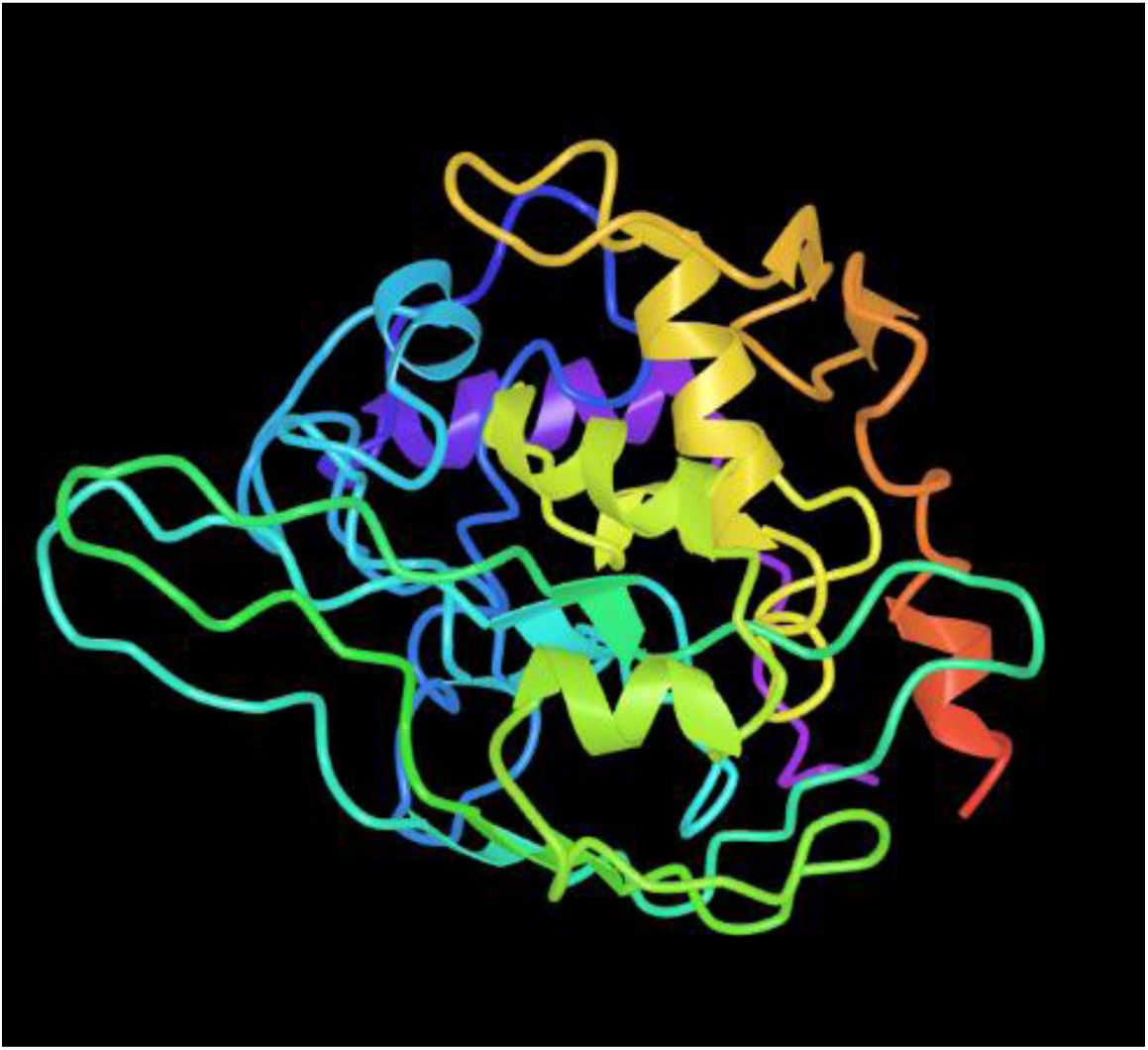
Predicted 3D structure of the vaccine construct, *Ras*IC-01vusing OSCR™.ai platform

**Figure 5.**
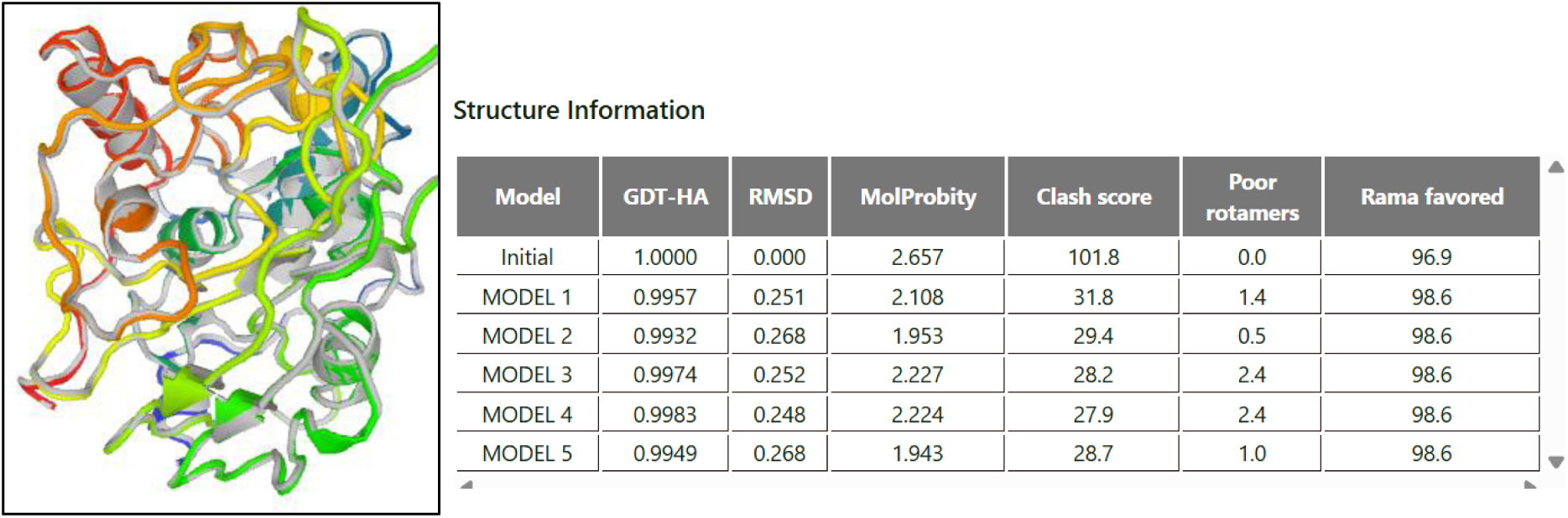
Resulting five models post-refinement with related structural information

**Figure 6.**
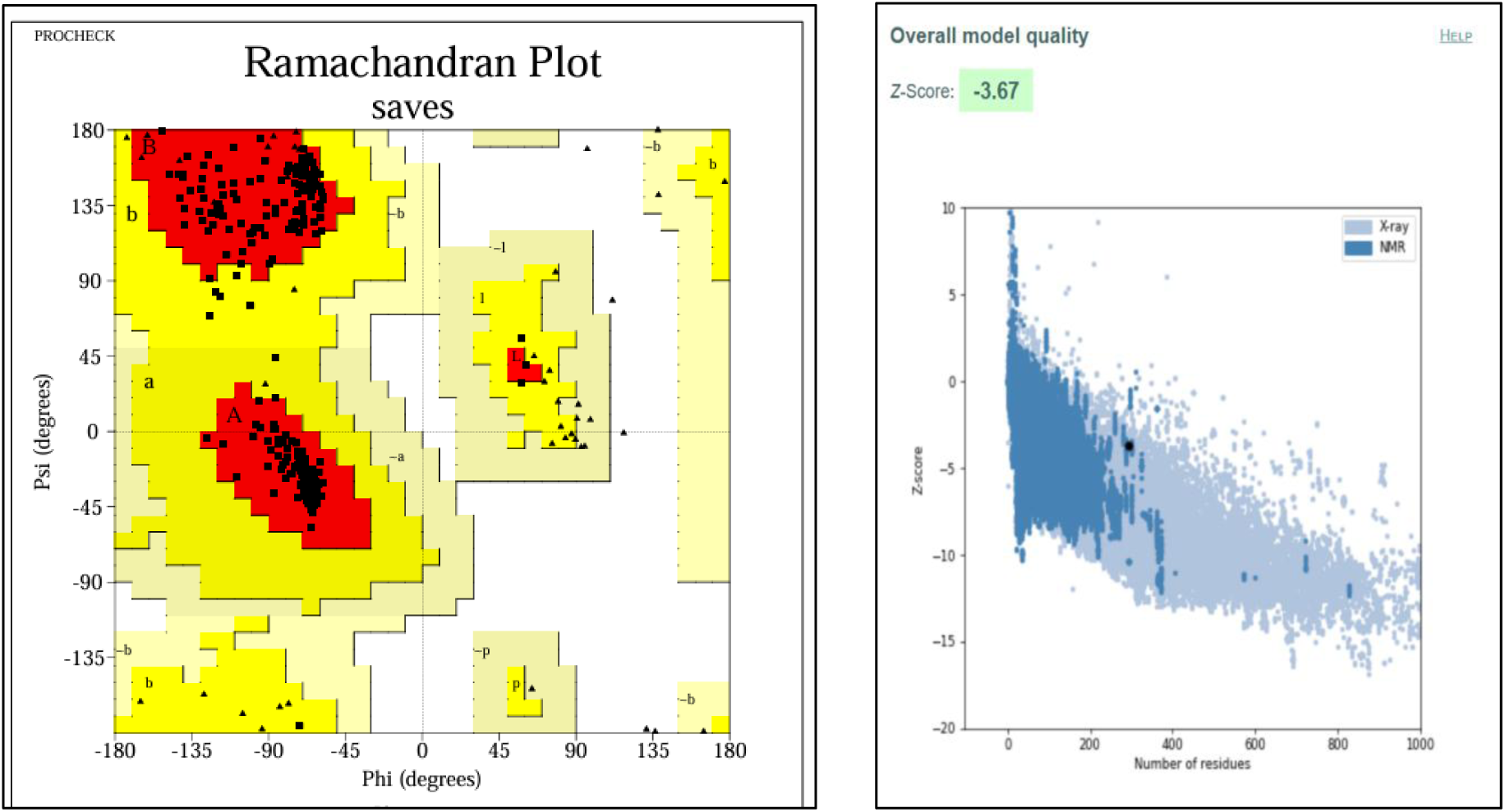
Ramachandran plot analysis showing 95.9% residues in most favoured region and ProSA-web score confirming the validity of the structure with a −3.67 Z score.

### Tertiary Structure Validation

The refined 3D structure was subjected to Ramachandran plot analysis using PROCHECK program as implemented in SAVES v6.1. The plot showed 95.9% in most favoured region and 4.1% in additional allowed region with 0.0% in disallowed region The structural validity of the modelled vaccine protein was assessed using ProSA-web. The model obtained a Z-score of −3.67, which falls within the range typically observed for native structures of comparable size resolved by NMR spectroscopy. In the ProSA energy plot, most residues exhibited negative energy values, indicating a predominantly stable folding pattern with only minor localized fluctuations. The position of the model’s Z-score within the reference distribution plot confirmed that the predicted structure is consistent with experimentally determined protein structures stored in the PDB. These findings supported the overall reliability and stereochemical quality of the modelled vaccine construct.

### MD Simulations of RasIC-01v

As depicted in Figures 7A-B, and 8A-B, the vaccine construct protein structure underwent drastic conformational changes after a time period of 15 ns. A specific portion of the protein structure starting from Ala274 up to Glu293 extended outward, thus adopting an open conformation. However, there were two other regions such as the one starting from Ser17 up to Gln30 and from Asp46 up to Pro73, which showed an RMS fluctuation in the range of 0.5-1.5 nm. With reference to Figure 8C, SASA analysis per residue complemented the RMS fluctuation analysis (Figure 8A). Residues Asp1, Asn3, Lys6, Pro9, Phe13, Glu20, Glu21, Lys23, Glu24, Gln30, Asp33, Ala45, Pro52, Pro54, Glu58, Val61, Val62, Pro68, Ser82, Glu101, Pro121, Pro124, Leu128, Arg131, Arg140, Ile141, Pro146, Glu148, Pro164, Pro166, Arg168, Leu171, Arg174, Pro180, Glu186, Leu191,Arg200, Trp204, Arg206, Lys226, Val232 Trp239, Pro250, Glu262, Glu272, Lys275, Asn278, Lys281, Phe287, Glu293 had their SASA in the range of 1-2.27 Nm³. Out of all the residues, Glutamate being polar has the highest SASA throughout the 100 ns trajectory; however surprisingly Proline also appears to have high SASA, which can be explained by the fact that the buried hydrophobic residues were exposed due to the opened/exposed conformation adopted by *Ras*IC-01v after 15 ns. Figures 8D and 8E validated the eminent unfolding of *Ras*IC-01v, as SASA curve (Figure 8D) took a sharp increment after 30 ns, which could be supported by Figure 8E, wherein the radius of gyration increased after 30 ns.

**Figure 7.**
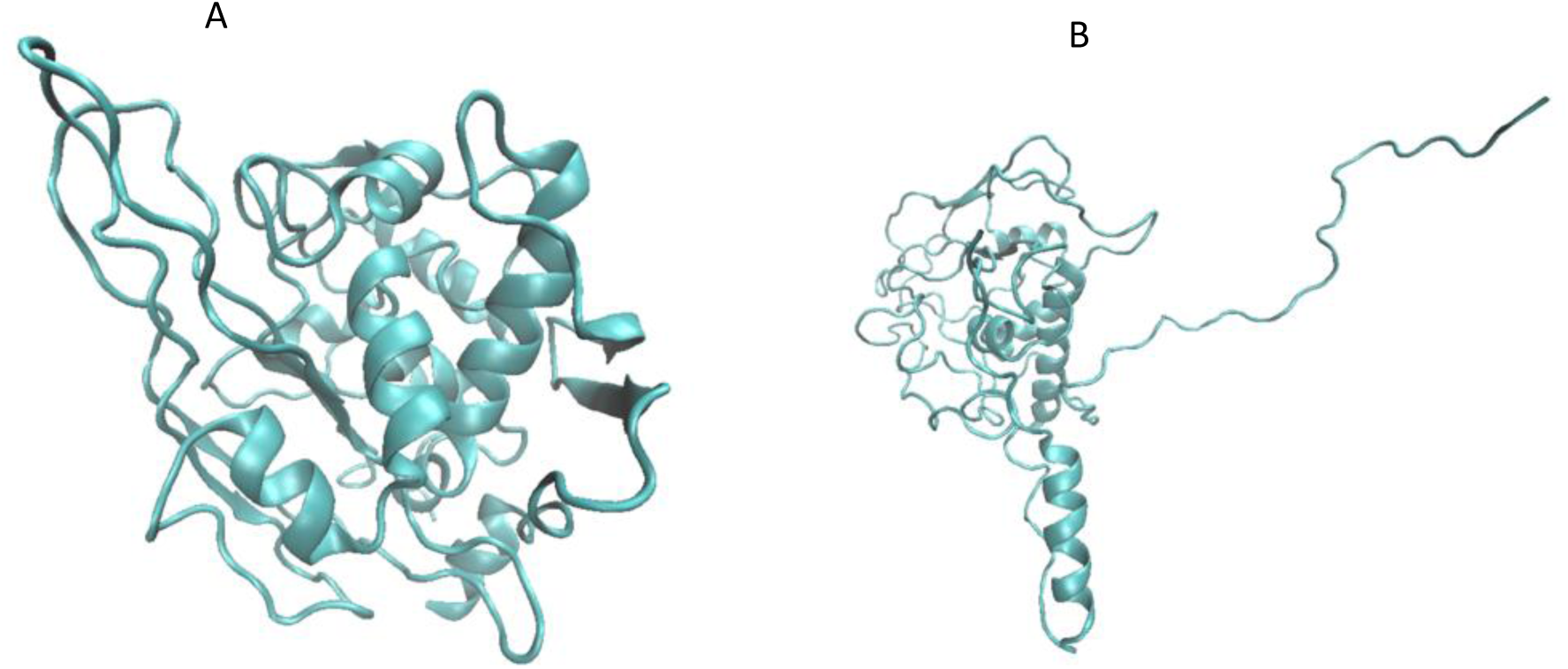
Comparison between initial predicted structure of RasIC-01v (A) and structure extracted from the last frame (100ns) of its solo simulation (B)

**Figure 8.**
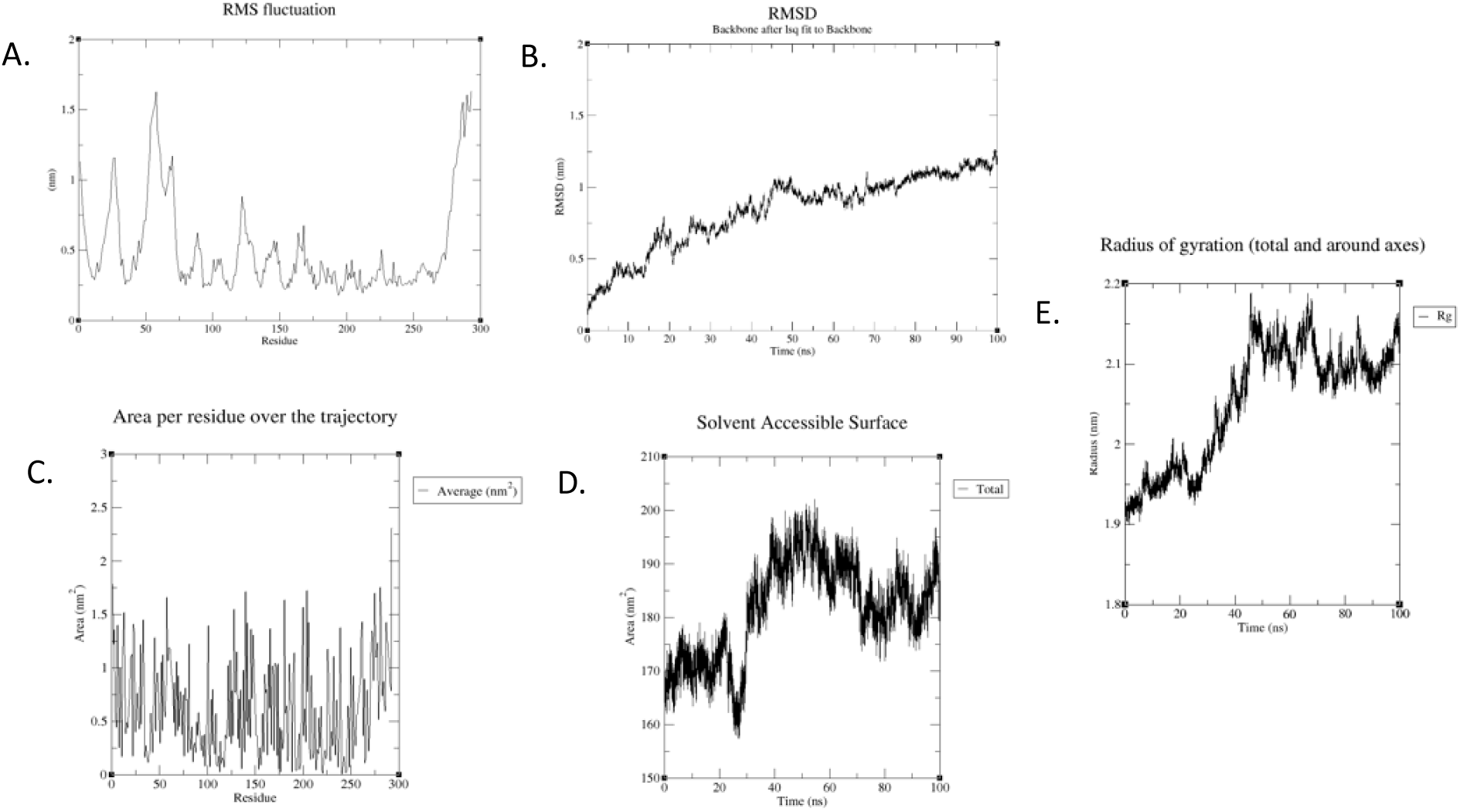
Post-simulation analysis for *Ras*IC-01v solo simulation. A. RMS Fluctuation analysis for each residue, B. RMSD vs time analysis for 100 ns trajectory, C. Solvent-accessible surface area (SASA) for each residue, D. SASA vs time analysis for 100 ns trajectory, E. Radius of gyration vs time analysis for 100 ns trajectory

With respect to the changes in the secondary structure (Figure 9A-B), most of the α-helix portion (highlighted in orange colour) showed significant RMS fluctuation, β-sheet portions (highlighted in red colour) showed moderate RMS fluctuation and 3_10_-helix portion (highlighted in yellow portion) showed the least RMS fluctuation, thus complementing the fact that β sheets were more stable than α helices.

**Figure 9.**
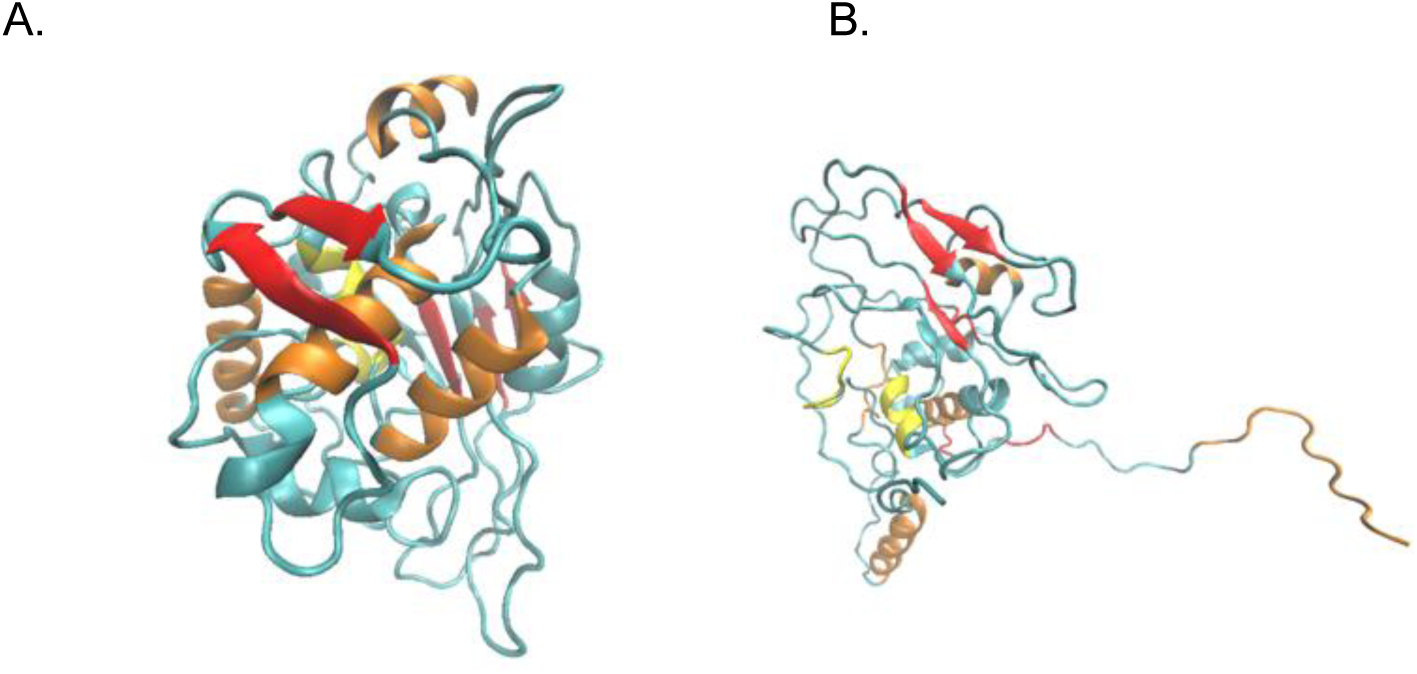
Comparison between the α-helices and β-sheets of initial predicted structure of Ras*IC-01v* and structure extracted from last frame (100 ns) of its solo simulation (B) depicted in different colours

### Docking of RasIC-01v (obtained after simulation) with TLR3

Docking runs performed using the *Ras*IC-01v structure obtained from the final frame of the standalone simulation produced an HDOCK score of −248.55. There were lesser interactions as opposed to the interactions obtained post simulation of GBM vaccine docked with TLR3 (Figures 10A-B). This outcome was attributable to the initially unfolded conformation selected for the docking protocol. As the docked pose stabilized throughout successive stages of the simulation, intermolecular interactions increased, resulting in stronger and more stable binding.

**Figure 10.**
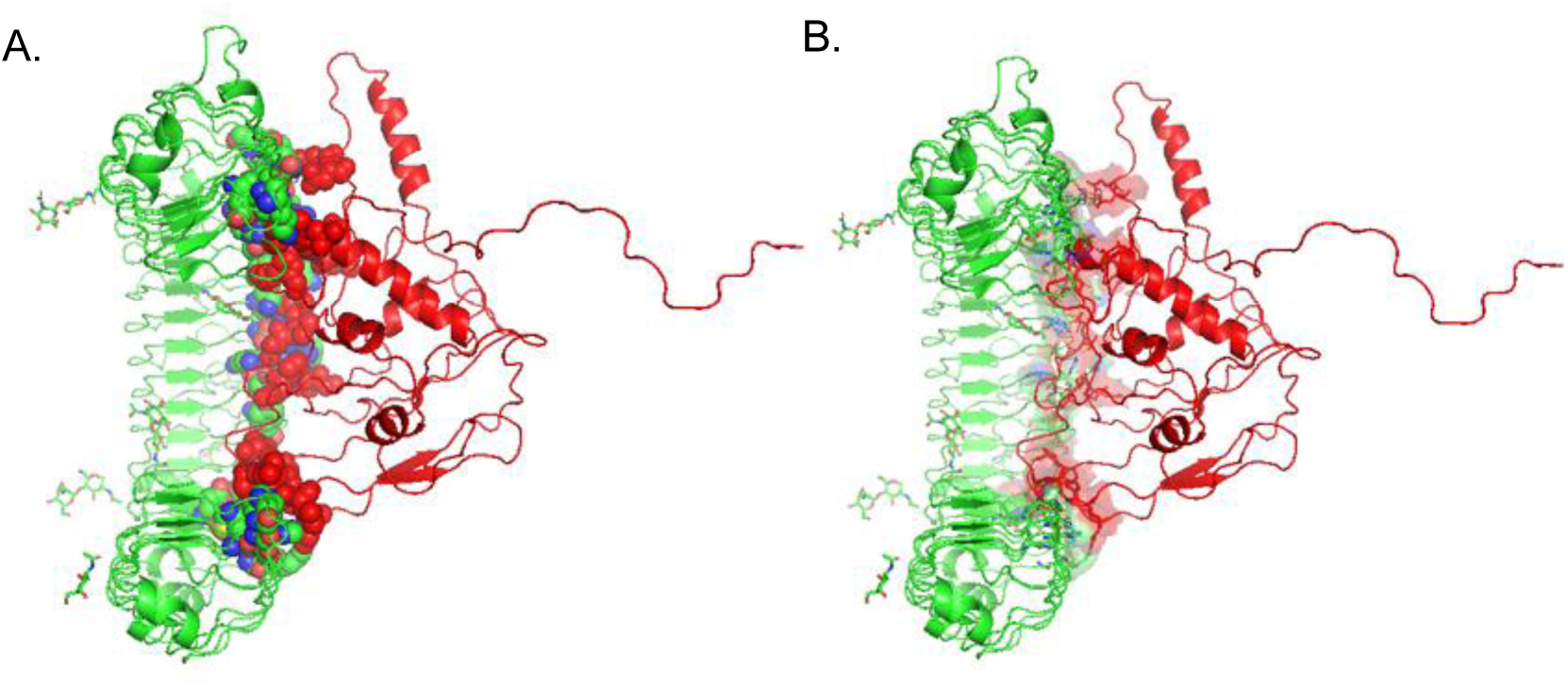
Alterations in *Ras*IC-01v secondary structure. A. Interface residues highlighted as spheres post docking, B. Interface residues highlighted as licorice.

Molecular docking analysis of the designed multi-epitope vaccine construct (*Ras*IC-01v) with the innate immune receptor TLR3 revealed the formation of multiple H-bonds at the receptor–ligand interface, indicating favourable initial recognition and binding orientation prior to molecular dynamics simulations (Figure 11A-F). Six H-bond interactions were observed in the docked complex: His127 (TLR3)–Pro124 (*Ras*IC-01v), Arg454 (TLR3)–Asp60 (*Ras*IC-01v), Ser103 (TLR3)–Arg168 (*Ras*IC-01v), Asn353 (TLR3)–Thr63 (*Ras*IC-01v), Asn482 (TLR3)–Asp33 (*Ras*IC-01v), and Tyr278 (TLR3)–Pro54 (*Ras*IC-01v). These interactions were distributed across multiple regions of the TLR3 binding surface, suggesting that the vaccine con-struct engaged a broad interface rather than a single localized contact site.

**Figure 11.**
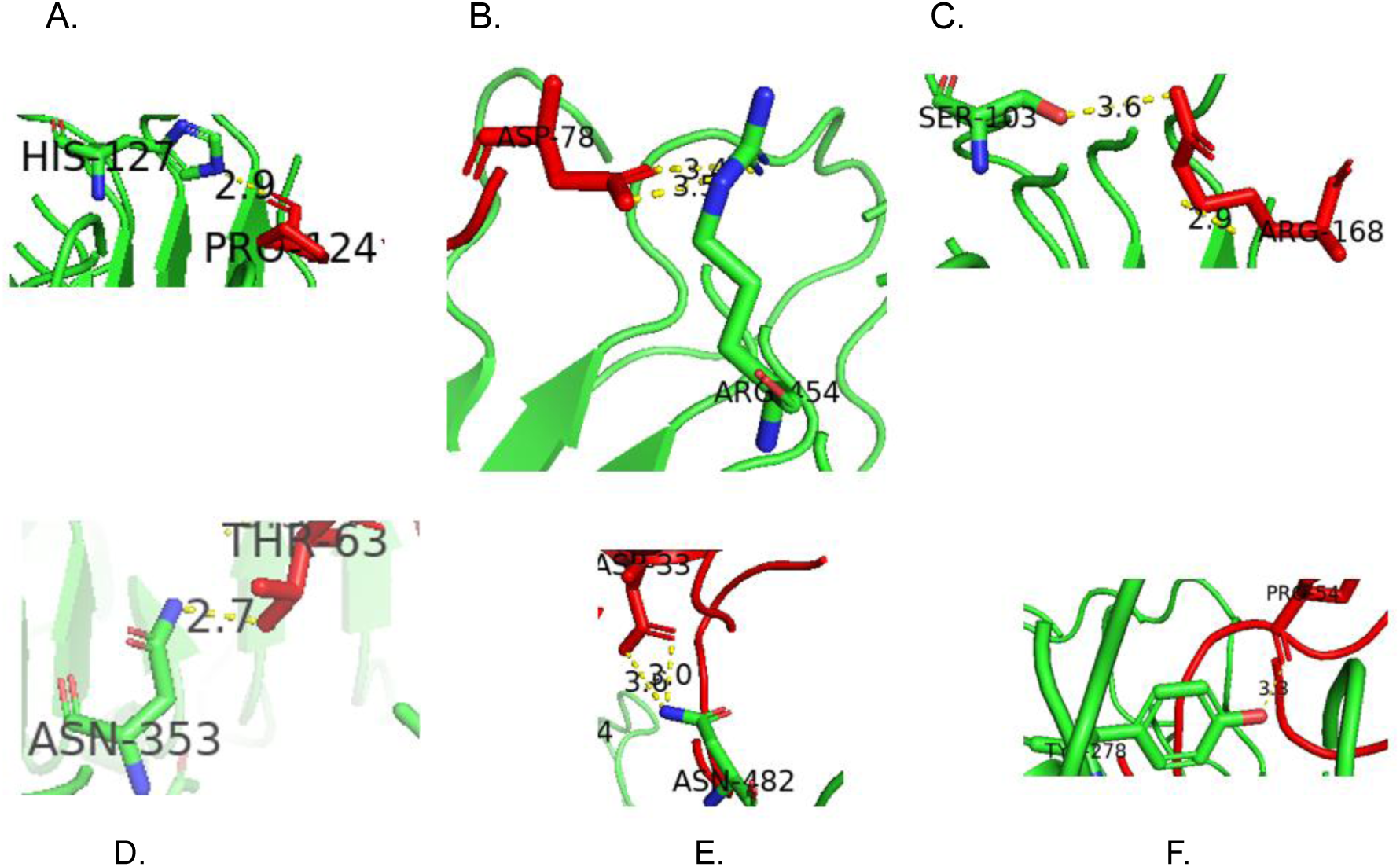
H-Bond formation among interface residues. A. H-bond between His127 (TLR3) and Pro124 (*Ras*IC-01v), B. H-bond between Arg454 (TLR3) and Asp60 (*Ras*IC-01v), C. H-bond between Ser103 (TLR3) and Arg168 (*Ras*IC-01v), D. H-bond be-tween Asn353 (TLR3) and Thr63 (*Ras*IC-01v), E. H-bond between Asn482 (TLR3) and Asp33 (*Ras*IC-01v), F. H-bond between Tyr278 (TLR3) and Pro54 (*Ras*IC-01v)

Several contacts involved complementary charged and polar residues (e.g., Arg–Asp and Asn–Asp), indicating electrostatic compatibility between the construct and receptor surface. Hydrogen bonds involving backbone atoms and structurally rigid residues such as proline further supported an anchored binding orientation of the ligand within the receptor groove. The presence of multiple interaction points implies that the docked pose was structurally plausible and capable of maintaining receptor engagement upon further refinement. Importantly, these interactions were identified from the pre-simulation docked complex and therefore represent the initial binding feasibility rather than the final stabilized complex. Nevertheless, the observed H-bond network suggested that the designed construct possessed appropriate spatial and physicochemical complementarity for TLR3 recognition, supporting its potential to initiate receptor-mediated immune activation in subsequent structural refinement and dynamic stability analyses.

### MD Simulations of the Docked Complex

Post-simulation trajectory analysis of the TLR3-*Ras*IC-01v complex demonstrated the formation of a well-defined and persistent binding interface (Figure 12). Extraction of interface residues from the equilibrated trajectory revealed a compact contact surface between the receptor and the vaccine construct, indicating that the initially docked orientation evolved into a structurally coherent interaction during molecular dynamics refinement. In the sphere representation (Figure 12A), interface residues from TLR3 (green) and *Ras*IC-01v (cyan) form a contiguous interaction patch rather than isolated point contacts. This clustering suggested co-operative stabilization, where multiple neighbouring residues collectively contributed to binding stability. The distribution of contacts across the concave ligand-recognition surface of TLR3 indicated that the construct occupied a receptor-compatible region known to participate in ligand engagement, supporting the structural plausibility of receptor activation.

**Figure 12.**
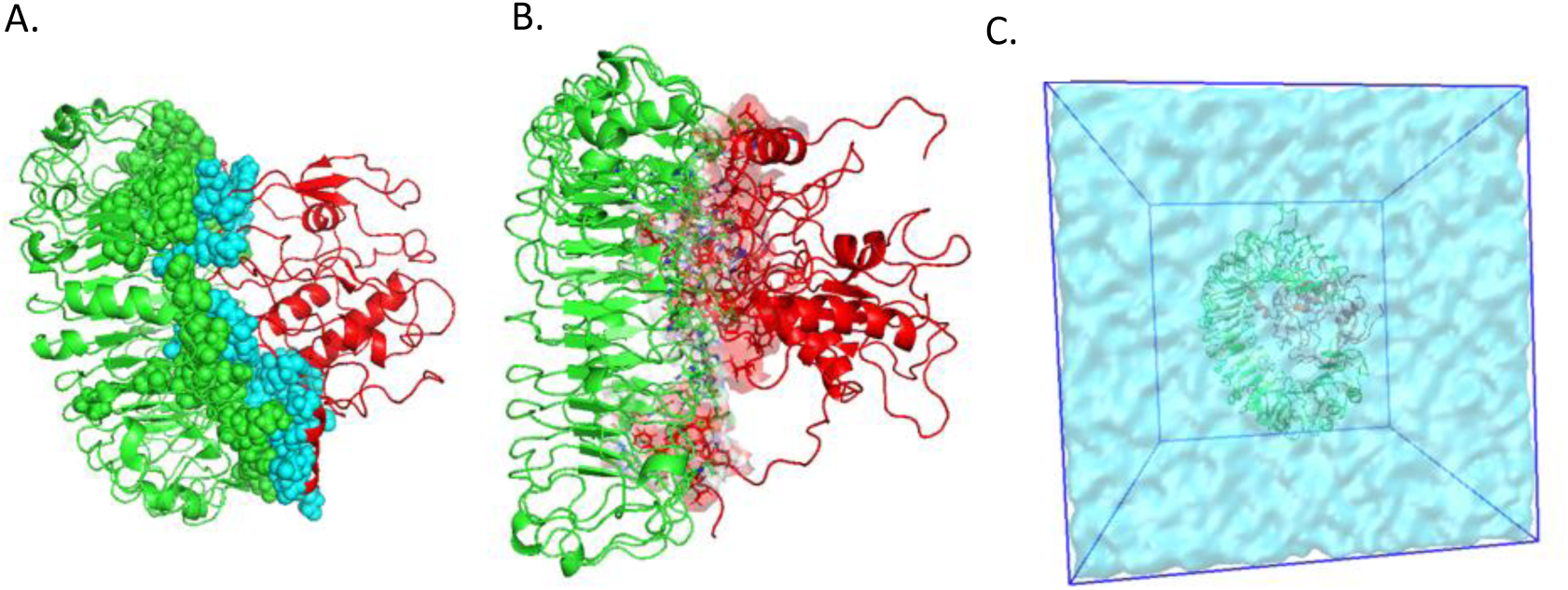
Interface residues extracted from post-simulation analyses. A. Interface residues represented as spheres (Green: TLR3, Cyan: *Ras*IC-01v), B. Interface residues represented as licorice, C. Complex solvated in the simulation box.

**Figure 13.**
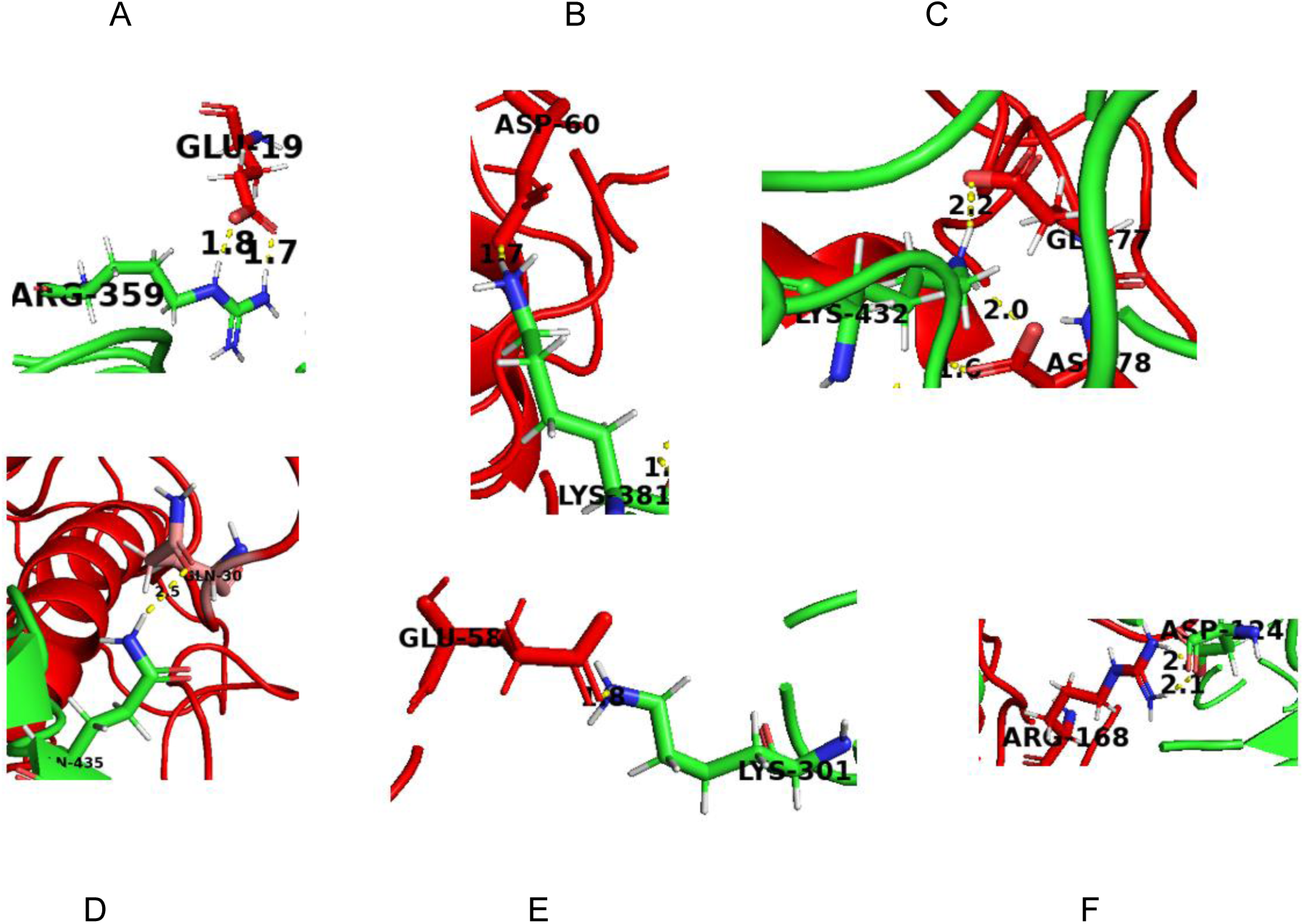
Interface residues forming H-bonds in the last frame of simulation. A. H-bond between Arg359 (TLR3) and Glu19 (*Ras*IC-01v), B. H-bond between Lys381 (TLR3) and Asp60 (*Ras*IC-01v), C. H-bond between Lys432 (TLR3) and Glu77 (*Ras*IC-01v) and Asp78 (*Ras*IC-01v), D. H-bond between Asn435 (TLR3) and Gln30 (*Ras*IC-01v), E. H-bond between Lys301 (TLR3) and Glu58 (*Ras*IC-01v), and F. H-bond between Asp124 (TLR3) and Arg168 (*Ras*IC-01v) Analysis of the final frame of the MD trajectory revealed a stabilized interaction network between the *Ras*IC-01v vaccine construct and TLR3, characterized by multiple persistent H-bonds across the receptor–ligand interface (Figure 13A-F). Compared with the pre-simulation docked pose, the equilibrated complex exhibited a reorganization of contacts, indicating conformational adaptation and strengthening of intermolecular interactions during simulation. Six principal H-bond interactions were identified: Arg359 (TLR3)-Glu19 (*Ras*IC-01v), Lys381 (TLR3)-Asp60 (*Ras*IC-01v), Lys432 (TLR3)-Glu77/Asp78 (*Ras*IC-01v), Asn435 (TLR3)-Gln30 (*Ras*IC-01v), Lys301 (TLR3)-Glu58 (*Ras*IC-01v), and Asp124 (TLR3)-Arg168 (*Ras*IC-01v).

Licorice representation of the same residues (Figure 12B) highlighted sidechain interdigitation between the receptor and the construct. The interaction network included polar, charged, and hydrophobic residues positioned in complementary orientations, indicating a combination of H-bonding, electrostatic interactions, and van der Waals contacts. Such multipoint anchoring is characteristic of stable peptide-receptor complexes and suggests reduced likelihood of spontaneous dissociation under physiological conditions. Solvation of the complex within the explicit water box (Figure 12C) confirmed proper system equilibration and absence of steric clashes or structural distortion after simulation. The complex remained centrally positioned within the periodic boundary box, indicating stable conformational behavior without drift or unfolding of the multi-epitope construct. The maintenance of the interaction interface in an aqueous environment further supported the thermodynamic feasibility of the complex.

Overall, post-simulation interface analysis indicated that *Ras*IC-01v established a stable and cooperative interaction surface with TLR3. The persistence of clustered interface residues and complementary sidechain packing suggested that the designed construct could maintain receptor engagement under dynamic conditions, strengthening its potential to function as an effective innate immune activator.

Notably, several of these contacts involved complementary charged residues (Lys/Glu, Arg/Glu, Lys/Asp, Asp/Arg), demonstrating strong electrostatic pairing at the interface. Such charge-assisted H-bonds are known to significantly enhance binding stability and specificity in protein–protein recognition.

The presence of multiple lysine-mediated interactions on the TLR3 surface suggested that positively charged receptor patches act as anchoring regions for negatively charged segments of the vaccine construct. This electrostatic complementarity likely contributed to maintaining ligand orientation and reducing conformational freedom after equilibration. Additionally, the dual interaction of Lys432 with both Glu77 and Asp78 indicated cooperative stabilization through a local interaction cluster rather than isolated contacts. Importantly, the persistence of these H-bonds in the last simulation frame confirms that the initially flexible multi-epitope construct underwent conformational accommodation to achieve a thermodynamically-favourable binding configuration. The shift toward a charge-complementary interaction network suggested induced-fit stabilization and improved interface packing compared with the initial docking structure.

Overall, the stabilized H-bond network observed after simulation supported robust receptor engagement and indicated that *Ras*IC-01v could maintain a stable association with TLR3 under dynamic conditions. This strengthened interaction pattern was consistent with the requirements for receptor recognition and supported the potential of the designed construct to effectively stimulate innate immune signalling.

Further inspection of the final simulation frame revealed additional stabilizing H-bond interactions at the TLR3-*Ras*IC-01v interface (Figure 14A-F), reinforcing the robustness of the receptor-ligand complex. Newly identified contacts included a cooperative interaction where Asn353 and Asn382 of TLR3 simultaneously engaged Arg79 of *Ras*IC-01v, indicating localized binding reinforcement through multi-residue anchoring. Additional polar interactions were observed between Thr629 (TLR3) and Lys281 (*Ras*IC-01v), Gln307 (TLR3) and Ala11 (*Ras*IC-01v), and Asp7 (TLR3) and Ser285 (*Ras*IC-01v), demonstrating that both terminal and internal segments of the construct contribute to receptor engagement. Notably, Lys432 and Lys383 of TLR3 form charge-assisted H-bonds with acidic residues Glu77 and Asp78 of the vaccine construct, establishing a recurrent electrostatic hotspot. The persistence of this negatively charged region interacting with multiple positively charged receptor residues suggested the formation of a stable electrostatic anchoring patch that likely governs binding orientation and retention. These additional interactions complemented the primary H-bond network identified previously, indicating progressive interface consolidation during simulation. The involvement of residues distributed across different regions of both TLR3, and the construct implied a broad contact surface rather than a single binding-pocket interaction. Such distributed binding is characteristic of productive receptor recognition and supports sustained receptor engagement under dynamic conditions.

**Figure 14.**
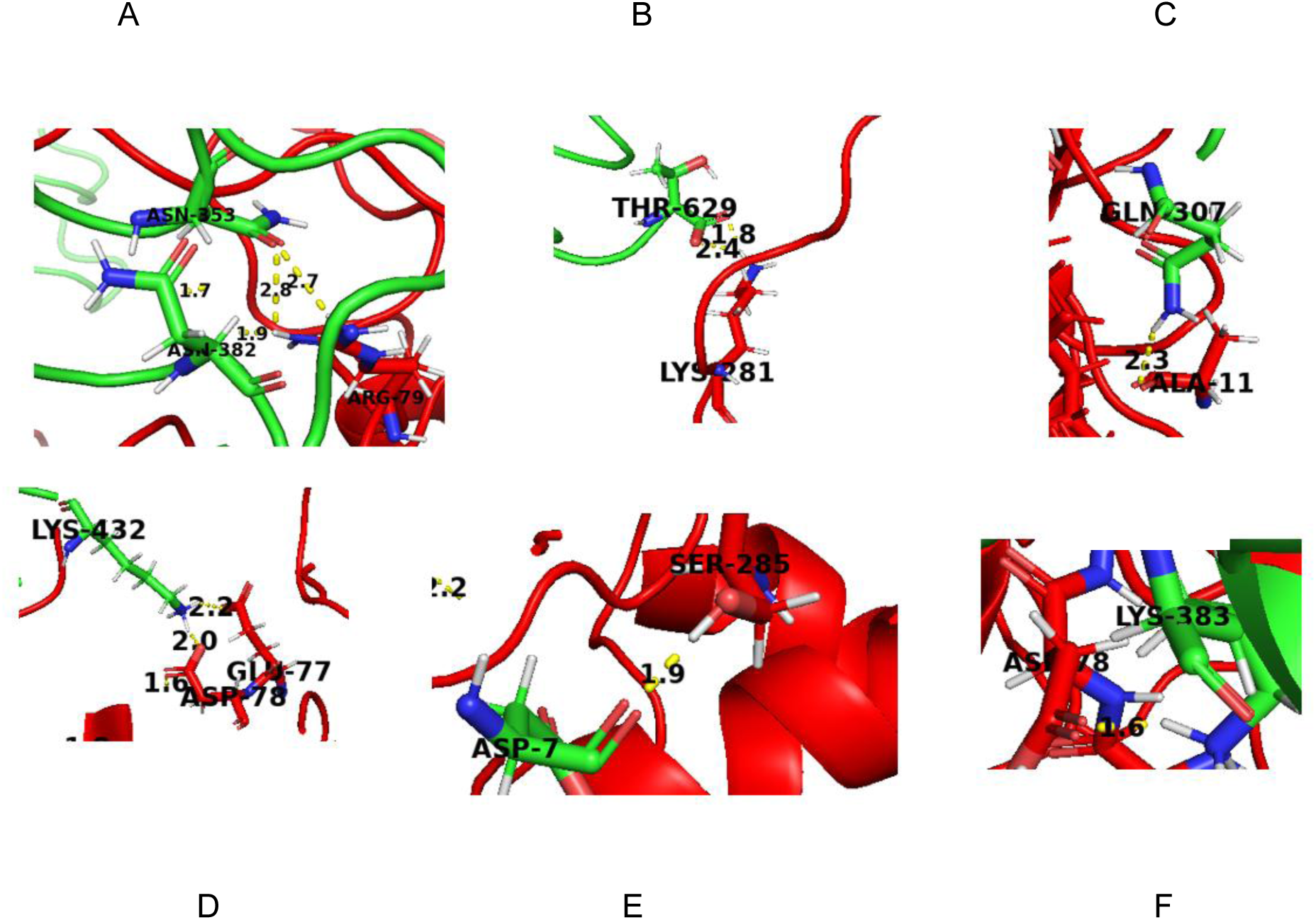
Interface residues forming H-bonds in the last frame of simulation. A. H-bond be tween Asn382 (TLR3), Asn353 (TLR3) and Arg79(*Ras*IC-01v), B. H-bond between Thr629 (TLR3) and Lys281 (*Ras*IC-01v), C. H-bond between Gln307 (TLR3) and Ala11 (*Ras*IC-01v), D. H-bond between Lys432 (TLR3) and Glu77 (*Ras*IC-01v), Asp78 (*Ras*IC-01v), E. H-bond between Asp7 (TLR3) and Ser285 (*Ras*IC-01v), F. H-bond between Lys 383 (TLR3) and Asp78 (*Ras*IC-01v)

Collectively, the expanded H-bond network confirmed enhanced structural stabilization of the TLR3-*Ras*IC-01v complex after equilibration, further supporting the capacity of the designed multi-epitope construct to maintain receptor association and potentially promote innate immune activation.

As shown in Figure 15A, the RMS fluctuation profile of the TLR3-*Ras*IC-01v complex exhibited a pronounced reduction in residue mobility compared with the vaccine-alone simulation (Figure 8A). Most residues displayed low fluctuation values, while elevated flexibility was confined primarily to terminal regions. The decrease in fluctuations across intermediate residues correlated with the formation of persistent H-bond interactions at the interface during the latter stages of the simulation, supporting stabilization of the bound conformation. The RMSD trajectory (Figure 15B) also showed improved structural stability relative to the solo vaccine system. An initial equilibration phase was followed by a decrease near ∼30 ns, indicating stabilization of the binding orientation. A transient increase between ∼45-65 ns corresponded to a conformational rearrangement involving residues Ala274-Glu293, which underwent a folding/closure transition. Following this event, the RMSD decreased and converged, suggesting attainment of a stable folded state for the complex. This compaction was further supported by the radius of gyration profile (Figure 15E), which mirrored the RMSD trend and indicated structural tightening after the mid-simulation folding event. In contrast, the solvent-accessible surface area (Figure 15C-D) remained comparable to the vaccine-alone system, consistent with formation of a partially compact yet solvent-exposed complex rather than global collapse. Finally, MM/GBSA calculations yielded a binding free energy of −20.73 kJ/mol, confirming energetically favorable and stable association between *Ras*IC-01v and TLR3. Collectively, reduced residue mobility, RMSD convergence after conformational rearrangement, and favorable binding energetics demonstrated that receptor binding induced structural stabilization and compaction of the vaccine construct, supporting a robust interaction with TLR3. Figure 16 depicts the conformational changes before and after docking, followed by MD simulations in *Ras*IC-01v structure, as seen from rearranged secondary structural elements.

**Figure 15.**
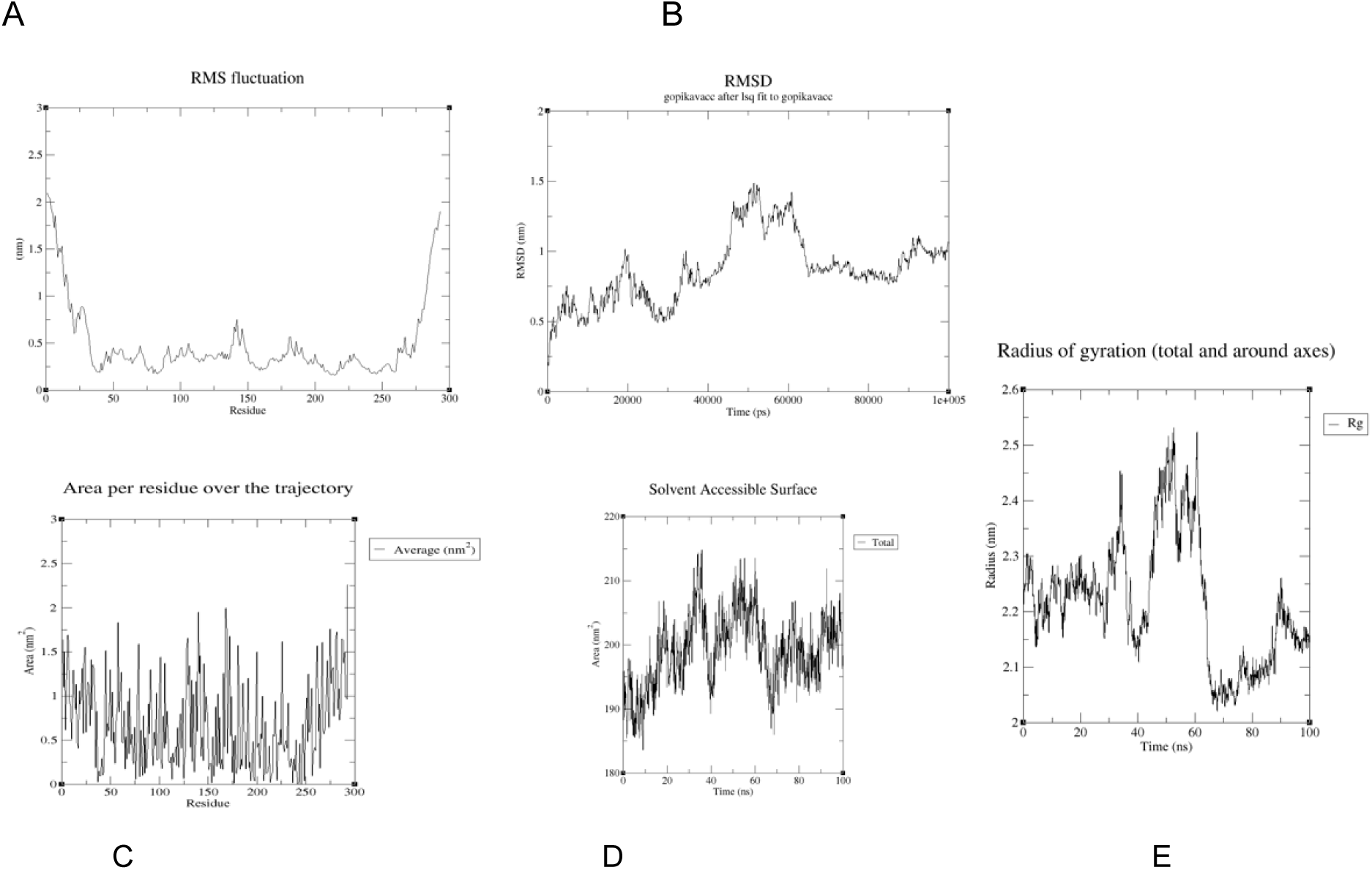
Post-simulation analyses of TLR3-*Ras*IC-01v complex. A. RMS Fluctuation analysis for each residue, B. RMSD vs time analysis for 100 ns trajectory, C. Solvent-accessible surface area for each residue, D. Solvent-accessible surface area vs time analysis for 100 ns trajectory, and E. Radius of gyration vs time analysis for 100 ns trajectory

**Figure 16.**
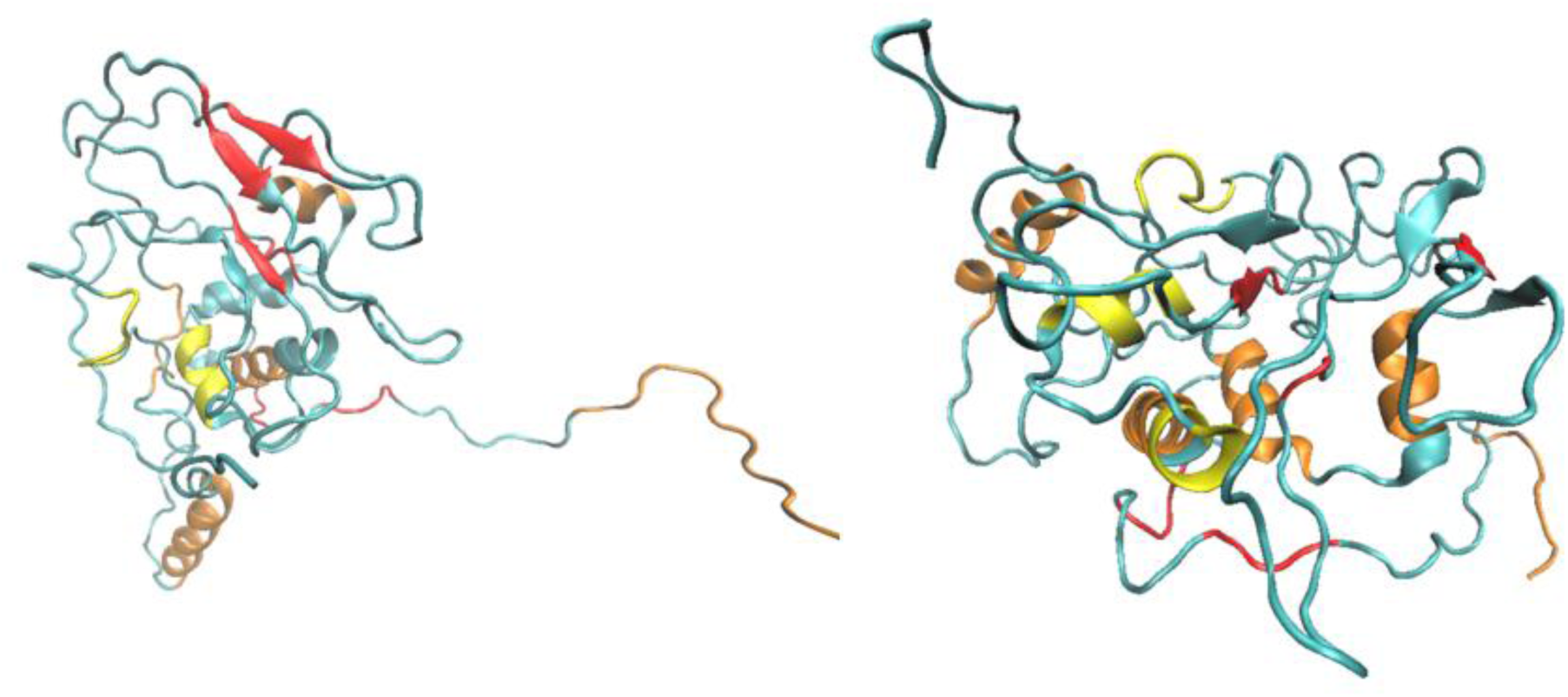
Conformational rearrangements in *Ras*IC-01v structure observed upon docking and subsequent MD simulation, illustrated through changes in secondary structural elements before and after simulation.

### Codon Optimization and In Silico Cloning

For optimizing the vaccine construct’s codon usage, VectorBuilder (https://en.vectorbuilder.com/tool/codon-optimization) tool was used for maximal protein expression in *Homo sapiens*. The optimized codon sequence had a length of 882 nucleotides (Figure 17) with a Codon Adaptation Index (CAI) score of 0.87 and GC % of 64.97, when *Homo sapiens* was used as host organism. The optimized sequence is as given below: >PDPN_Homo sapiens

**Figure 17.**
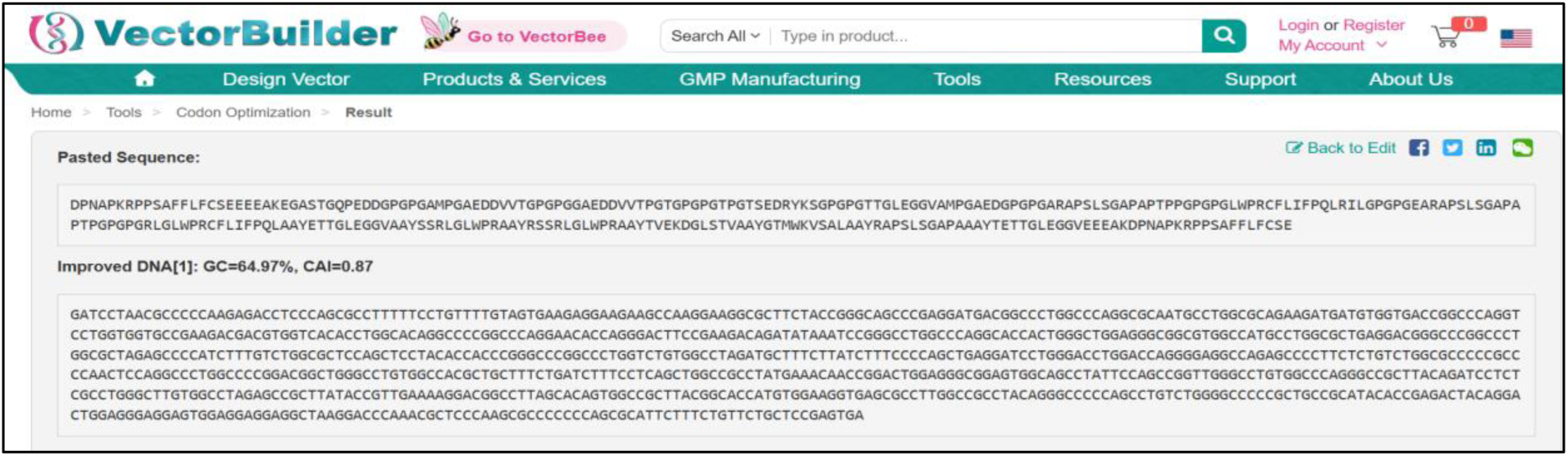
In silico codon optimization using VectorBuilder

**Figure 18.**
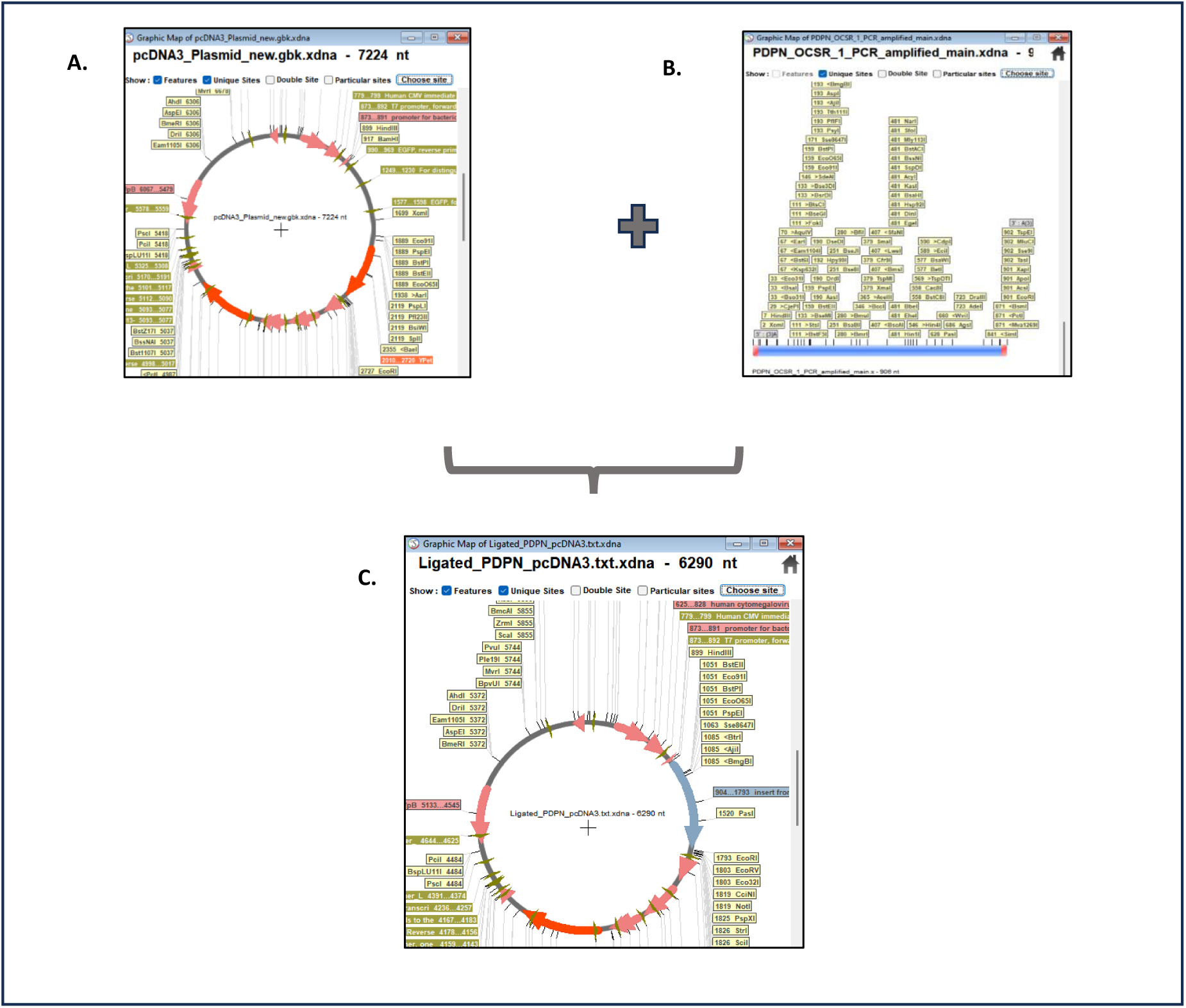
In silico cloning. A. pcDNA3 vector: with 5’-HindIII and 3’-EcoRI restriction sites, B. PDPN insert fragment, and C. Ligated product (shown in blue).

**Figure 19.**
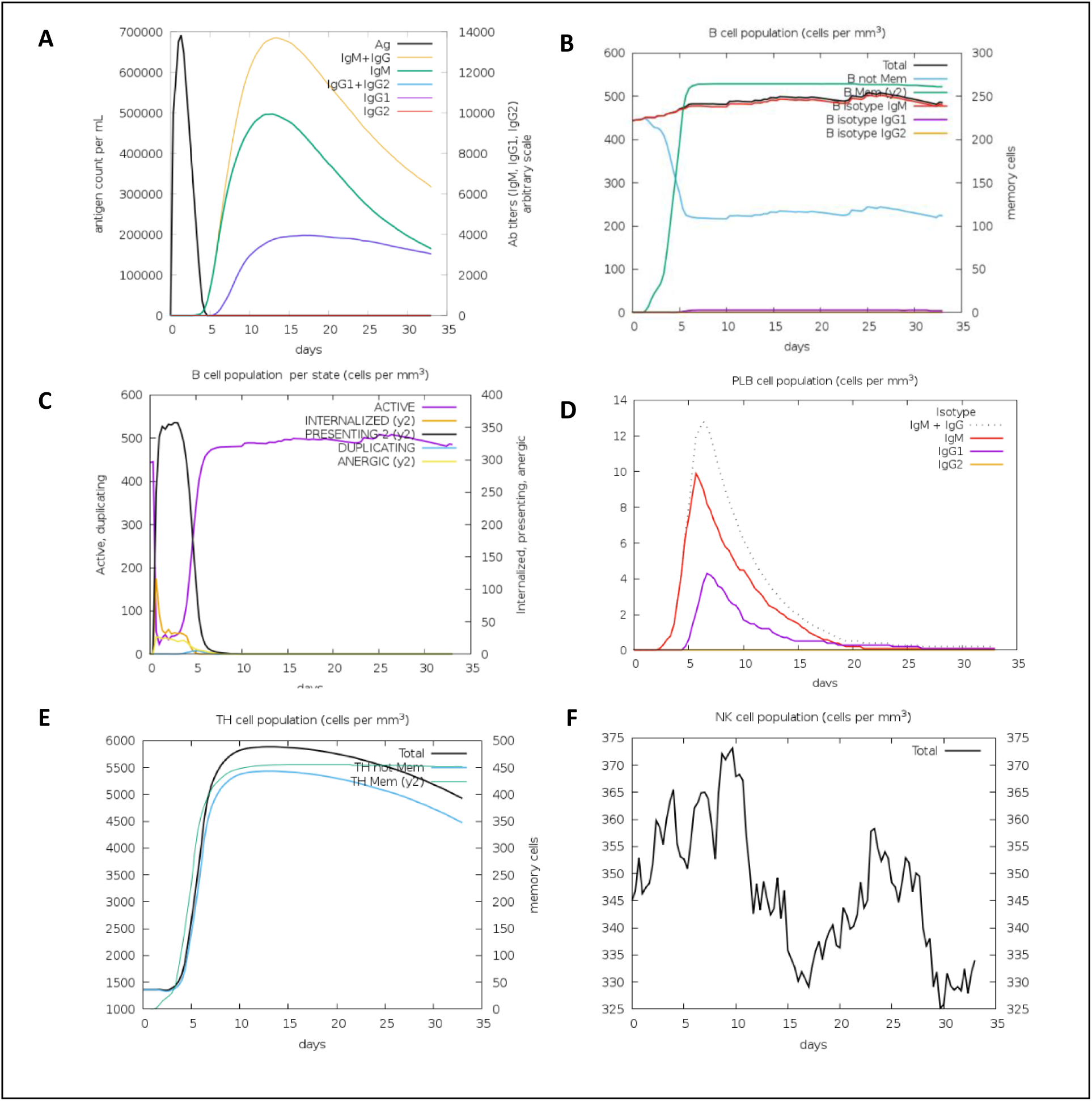
A. Antigen and antibody titres over time: Antigen levels peak early and decline, while IgM rises first and IgG1 becomes the dominant sustained antibody response, with IgG2 remaining minimal, B. Total B-Cell and Memory B-Cell Response: Total and memory B cells expand rapidly after vaccination and remain stable, indicating a sustained humoral response, C. B-Cell Functional States: Activated, presenting, and duplicating B-cell states appear early and contract as the response stabilizes, while anergic cells remain low, D. Plasma B-Cell (PLB) Dynamics: Plasma B-cell isotypes show a strong early IgM and IgG1 secretion peak, with IgG2 production staying very limited, E. Helper T-Cell (TH) Response: Helper T cells expand quickly and plateau, with memory TH subsets persisting throughout the simulation, F. Natural Killer (NK) Cell Activity: NK cells show an early activation fluctuation followed by a stable maintenance phase indicative of innate immune engagement.

GATCCTAACGCCCCCAAGAGACCTCCCAGCGCCTTTTTCCTGTTTTGTAGTGAA-GAGGAA-GAAGCCAAGGAAGGCGCTTCTACCGGGCAGCCCGAGGATGACGGCCCTGGCCCA GGCGCAATGCCTGGCGCAGAAGATGATGTGGTGAC-CGGCCCAGGTCCTGGTGGTGCCGAA-GACGACGTGGTCACACCTGGCACAGGCCCCGGCCCAGGAACACCAGGGACTTCC GAAGACAGATATAAATCCGGGCCTGGCCCAGGCACCACTGGGCTG-GAGGGCGGCGTGGCCATGCCTGGCGCTGAGGACGGGCCCGGCCCTGGCGCTAGA GCCCCATCTTTGTCTGGCGCTCCAGCTCCTACAC-CACCCGGGCCCGGCCCTGGTCTGTGGCCTAGATGCTTTCTTATCTTTCCCCAGCTG AGGATCCTGGGACCTGGACCAGGG-GAGGCCAGAGCCCCTTCTCTGTCTGGCGCCCCCGCCCCAACTCCAGGCCCTGGCC CCGGACGGCTGGGCCTGTGGCCAC-GCTGCTTTCTGATCTTTCCTCAGCTGGCCGCC-TATGAAACAACCGGACTGGAGGGCGGAGTGGCAGCCTATTCCAGCCGGTTGGGCC TGTGGCCCAGGGCCGCTTACAGATCCTCTCGCCTGGGCTTGTGGCCTA-GAGCCGCTTATAC-CGTTGAAAAGGACGGCCTTAGCACAGTGGCCGCTTACGGCACCATGTGGAAGGTG

AGCGCCTTGGCCGCCTACAGGGCCCCCAGCCTGTCTGGGGCCCCCGCTGCCG-CATACACCGA-GACTACAGGACTGGAGGGAGGAGTGGAGGAGGAGGCTAAGGACCCAAACGCTCC CAAGCGCCCCCCCAGCGCATTCTTTCTGTTCTGCTCCGAGTGA

The optimized codon was used as fragment for in silico PCR. Forward and reverse primers were designed to check if the fragment was getting amplified. Forward primer with sequence, 5’ GCCACCAAGCTTATGGATCCTAACGCCCCC 3’ had Tm of 51.6 °C for annealing template region and the reverse primer with sequence, 5’ GAATTCTAATCACTCGGAGCAGAA 3’, had Tm of 46.1°C for the same. The PDPN optimized codon got amplified and gave a 906 nt long fragment after amplification. Vector pcDNA3 was chosen since it is usually used as mammalian expression vector in CHO cells. The vector GenBank sequence was downloaded from addgene (https://www.addgene.org/64927/). Vector was 7225bp long with an already present insert named TORCAR tagged by 5’ HindIII and 3’ EcoRI. Restriction digestion was used to remove the already present insert and then the PDPN PCR amplified fragment was ligated using SerialCloner software (http://serialbasics.free.fr/Serial_Cloner.html).

### Immune Simulation Analysis

Immune simulation showed a strong primary response characterized by a rapid rise in IgM followed by class switching to IgG, with IgG1 and IgG2 remaining dominant through the later phases. A marked increase in memory B-cell populations was observed after the secondary exposure, and subsequent antigen encounters generated faster and higher antibody responses, demonstrating effective immune recall. T-helper cell subsets, particularly Th1 and Th2, expanded early and stabilized over time, while cytotoxic T cells showed a gradual increase in their activated and memory fractions. The cytokine and interleukin profiles indicated sustained immune stimulation without excessive inflammatory peaks, suggesting a controlled and antigen-specific response. Overall, the simulation output supports the potential of the construct to induce long-lasting humoral and cellular immunity with efficient memory formation.

## Conclusions

The present work outlines the design and analysis of a multi-epitope subunit vaccine, *Ras*IC-01v, based on the target Podoplanin (PDPN) using an immunoinformatics approach for the treatment of glioblastoma, which enriches the resurfacing of this type of cancer and its related aggressive behaviour. PDPN expression analysis indicated high transcriptome expression of this target compared to the normal human brain, indicating the suitability of this target to design this anti-glioblastoma vaccine candidate. The B and T cell epitope regions were fused together to form the final recombinant structure with favourable antigenicity, stability, and safety parameters and were also indicated to form stable interactions with the innate immunity biomolecules using structural modelling and immune simulation analysis to provide both cellular and humoral immunity.

Despite the in silico results offering a compelling rationale for the proposed design of the vaccine, biological validation would be required to show its significance. Moving ahead, it would be important to validate the expression of PDPN in glioblastoma cell lines and tissue, followed by the cloning of the vaccine construct into an expression vector and subsequent expression in CHO cells. This secreted recombinant protein can then be isolated from cell culture supernatants and used to activate human peripheral blood mononuclear (PBMC) cells in vitro. Analysing interferon-γ production by ELISA assay will provide a direct measure of T-cell activation, as well as a first look at the potential immunogenicity of a vaccine based on this predictive model as a whole. This body of work establishes a sound computational paradigm for PDPN-targeted vaccine development.

## Abbreviations

APC: Antigen-Presenting Cell
AI: Artificial Intelligence
BBB: Blood–Brain Barrier
BIRC5: Baculoviral IAP Repeat Containing 5 (Survivin)
BLAST: Basic Local Alignment Search Tool
CAI: Codon Adaptation Index
CAR-T: Chimeric Antigen Receptor T-cell
CESC: Cervical Squamous Cell Carcinoma
CHO: Chinese Hamster Ovary
CTL: Cytotoxic T Lymphocyte
ELISA: Enzyme-Linked Immunosorbent Assay
GBM: Glioblastoma Multiforme
GEPIA: Gene Expression Profiling Interactive Analysis
GTEx: Genotype-Tissue Expression
HDOCK: Hybrid Docking Server
HLA: Human Leukocyte Antigen
HMGB1: High Mobility Group Box 1
HNSC: Head and Neck Squamous Cell Carcinoma
HPC: High-Performance Computing
HTL: Helper T Lymphocyte
IC_50_: Half Maximal Inhibitory Concentration
IEDB: Immune Epitope Database
Ig: Immunoglobulin
IL: Interleukin
LUSC: Lung Squamous Cell Carcinoma
MAST: Motif Alignment Search Tool
MD: Molecular Dynamics
MERCI: Motif Emerging and with Classes – Identification
MHC: Major Histocompatibility Complex
ML: Machine Learning
MMGBSA: Molecular Mechanics Generalized Born Surface Area
MMPBSA: Molecular Mechanics Poisson–Boltzmann Surface Area
MySQL: My Structured Query Language
NK: Natural Killer Cell
NMR: Nuclear Magnetic Resonance
NPT: Constant Number, Pressure and Temperature
NVT: Constant Number, Volume and Temperature
PAAD: Pancreatic Adenocarcinoma
PBMC: Peripheral Blood Mononuclear Cell
PCR: Polymerase Chain Reaction
PDPN: Podoplanin
PDB: Protein Data Bank
pLDDT: Predicted Local Distance Difference Test
PLB: Plasma B-Cell
ProSA: Protein Structure Analysis Server
RMSD: Root Mean Square Deviation
RNA: Ribonucleic Acid
RMSF: Root Mean Square Fluctuation
SASA: Solvent Accessible Surface Area
SKCM: Skin Cutaneous Melanoma
SOX: SRY-Related HMG-Box Transcription Factor
TAA: Tumor-Associated Antigen
TCGA: The Cancer Genome Atlas
TGCT: Testicular Germ Cell Tumours
THC: T-Helper Cell
THCA: Thyroid Carcinoma
TLR3: Toll-Like Receptor 3
TPM: Transcripts Per Million
VMD: Visual Molecular Dynamics

